# Lineages of embryonic stem cells show non-Markovian state transitions

**DOI:** 10.1101/2020.11.23.372268

**Authors:** Tee Udomlumleart, Sofia Hu, Salil Garg

## Abstract

Pluripotent embryonic stem cells (ESCs) contain the ability to constitute the cell types of the adult vertebrate through a series of developmental state transitions. In culture, ESCs reversibly transition between states in a manner previously described as stochastic. However, whether ESCs retain memory of their previous states or transition in a memoryless (Markovian) process remains relatively unknown. Here we show lineages of ESCs do not exhibit the Markovian property: their previous states and kin relations influence future choices. In a subset of lineages, related ESCs remain likely to occupy the same state weeks after labeling. Unexpectedly, the distribution of lineages across states away from the equilibrium point predicted by a Markov model remains consistent over time, suggesting a conservation of informational entropy in this system. Additionally, some lineages appear highly dynamic in their ability to switch states but do not dominate the culture, suggesting that state switching is a separable property from growth. Together, these data suggest ESC state transitions are a proscribed process governed by additional variables.

## Introduction

Stochastic processes have been described to play a role in multiple mammalian developmental pathways, ranging from hematopoiesis to fate choice of retinal progenitors (Boije et al., 2014; Till and Mc, 1961). For example, the development of mature retinal cell types from retinal precursor cells follows consistent probabilities as precursor cells choose a lineage fate without any apparent regard to environment or history, and therefore has been termed stochastic (Losick and Desplan, 2008). In probability theory, a stochastic process that does not exhibit memory of its history is termed a Markovian process and is said to possess the Markov property. Therefore, for a memoryless (Markovian) stochastic process, the probability of visiting each state next depends only on the current state and not any preceding states. However, few studies of biological processes termed stochastic have formally assessed whether these processes possess the Markov property. In biological development, cell states are often thought of as the expression of groups of genes at or near specific levels for each gene (Garg and Sharp, 2016). Knowing whether the history of a process influences future cell states is of particular interest for reversible transitions, where multiple paths could lead to the present state. Such reversible transitions occur in many contexts, such as maintenance of airway epithelium or intestinal crypts (de Sousa and de Sauvage, 2019; Nabhan et al., 2018; Pardo-Saganta et al., 2015; Tata et al., 2013; Tetteh et al., 2016), or in reprogramming experiments whereby differentiated cell types are induced to pluripotent cell states (Biddy et al., 2018). Understanding whether the history of prior states influences the probability of reaching particular future states will be important for understanding development and homeostasis of mammalian tissues.

One context in which to consider reversible state transitions is early embryogenesis in mammals, whereby loss of particular cells can lead to replacement through the developmental plasticity of neighbors (Chen et al., 2018; Martinez Arias et al., 2013). Embryonic stem cells (ESCs) provide an interesting model of early development, as these cells are derived from the inner mass of the blastocyst and can form all tissues of the adult vertebrate organism, and ESC state transitions in culture mimic developmental state transitions in embryos (Neagu et al., 2020; Shahbazi et al., 2017). ESCs show remarkable heterogeneity in the expression of key transcription factors, such as the pluripotency genes Nanog and Sox2 (Abranches et al., 2014; Chakraborty et al., 2020; Chambers et al., 2007; Filipczyk et al., 2015; Kalmar et al., 2009; Klein et al., 2015; Kumar et al., 2014; Singer et al., 2014; Ying et al., 2008), and heterogeneous expression in ESCs has been previously classified into discrete states with different developmental potential (Abranches et al., 2014; Filipczyk et al., 2015; Kalmar et al., 2009). ESCs dynamically interconvert between states, transitioning back and forth under standard culture conditions (Chakraborty et al., 2020; Filipczyk et al., 2015; Singer et al., 2014). Previous studies characterizing the dynamics of state transitions in this system have focused on states defined by levels of Nanog, and have utilized fluorescent reporters in addition to antibody staining or fluorescence in situ hybridization (Chambers et al., 2007; Filipczyk et al., 2015; Singer et al., 2014). These studies have described the process of interconversion between states as stochastic, using measurements typically taken over timescales on the order of hours (Abranches et al., 2014; Hormoz et al., 2016; Ochiai et al., 2014; Singer et al., 2014). However, whether or not ESCs possesses the Markov property has not been extensively evaluated, and ESC state transitions over longer timescales have not been explored. ESC state transitions in culture mimic developmental state transitions in embryos (Neagu et al., 2020; Shahbazi et al., 2017), and early embryogenesis in mammals can involve reversible transitions whereby loss of particular cells leads to replacement through the developmental plasticity of neighbors (Chen et al., 2018; Martinez Arias et al., 2013). Together, this makes ESCs a particularly interesting model system to consider memory of states due to their ability to generate a diverse array of cell fates and their exhibiting reversible state transitions in culture.

One method to assess whether state transitions are a Markovian process is to examine whether daughter and cousin cells within a lineage of ESCs maintain a relationship in the states they occupy. In a Markovian process, each cell makes a state choice independent of its history, so the correlation of cell states between kin cells relaxes over time. As a result, eventually all cell lineages converge towards a consistent distribution of cell states as mixing amongst states occurs independently within each lineage. That is, for a stochastic memoryless process all lineages should converge to the same distribution of states. Measuring how close or far a dynamical system is from this convergence point represents a type of informational entropy (Baez and Pollard, 2016). Whether or not ESC state transitions are Markovian processes and the degree to which they diverge from a Markovian model over long timescales is unknown.

Here we characterize the dynamics of ESC state transitions amongst three interconverting states defined by levels of Nanog and Sox2 that represent distinct gene expression programs related to development (Chakraborty et al., 2020). We genetically barcode ESCs, expand the population, and observe the proportion of each ESC lineage in each state over time. We find state history for each ESC lineage influences future state transitions, and therefore ESCs do not exhibit the Markovian property on the measured timescale. Surprisingly, a subset of ESC lineages shows apparent concerted state transitions weeks after the barcode label is applied. These lineages exhibit a high frequency of concerted state transitions but remain a consistent proportion of the total population. This subset of lineages shows small but significant correlation in its degree of transition between replicate experiments. Finally, we measure the distribution of all lineages across state space as compared to the predictions of a Markovian model and quantify the difference as a type of informational entropy we term lineage entropy. Strikingly, the distribution of lineage entropy appears conserved over time. Together these data show ESC state transitions do not possess the Markov property and raise the question of whether this process is stochastic or governed by additional variables.

## Results

### Generation and tracking of ESC lineages over time

To assess the dynamics of ESC state transitions over time, we constructed an ESC reporter line compatible with barcoding and state readout. We generated ESC with heterozygous insertions of fluorophore tags at the endogenous loci of *Nanog* and *Sox2* (*GFP*-P2A-*Nanog* and *Sox2*-P2A-*mCerulean3* respectively, Fig. S1A). We previously divided these cells into three predominant states of Nanog and Sox2 expression (State 1 = High Nanog and High Sox2, State 2 = Low Nanog and High Sox2, and State 3 = Low Nanog and Low Sox2; Fig. 1A and (Chakraborty et al., 2020)) in ESC. We transduced ESCs with a previously described lentiviral barcoding vector (Bhang et al., 2015) at a low multiplicity of infection, ensuring each cell received <= 1 barcode (Fig. S1). After selecting for ~100,000 transduced, labeled cells representing at least 5,341 distinct barcoding events we expanded the population to 10^8^ total cells, allowing each barcoded ESC the chance to expand to an estimated ~15,000 cells distributed across all three ESC states, and cultured cells together (Fig. 1B). We refer to these expanded, single ESC derived cells as ESC lineages since the incorporated lentiviral barcode will be copied in each progeny cell, marking all ESC with the same unique barcode as kin.

**Figure 1.**
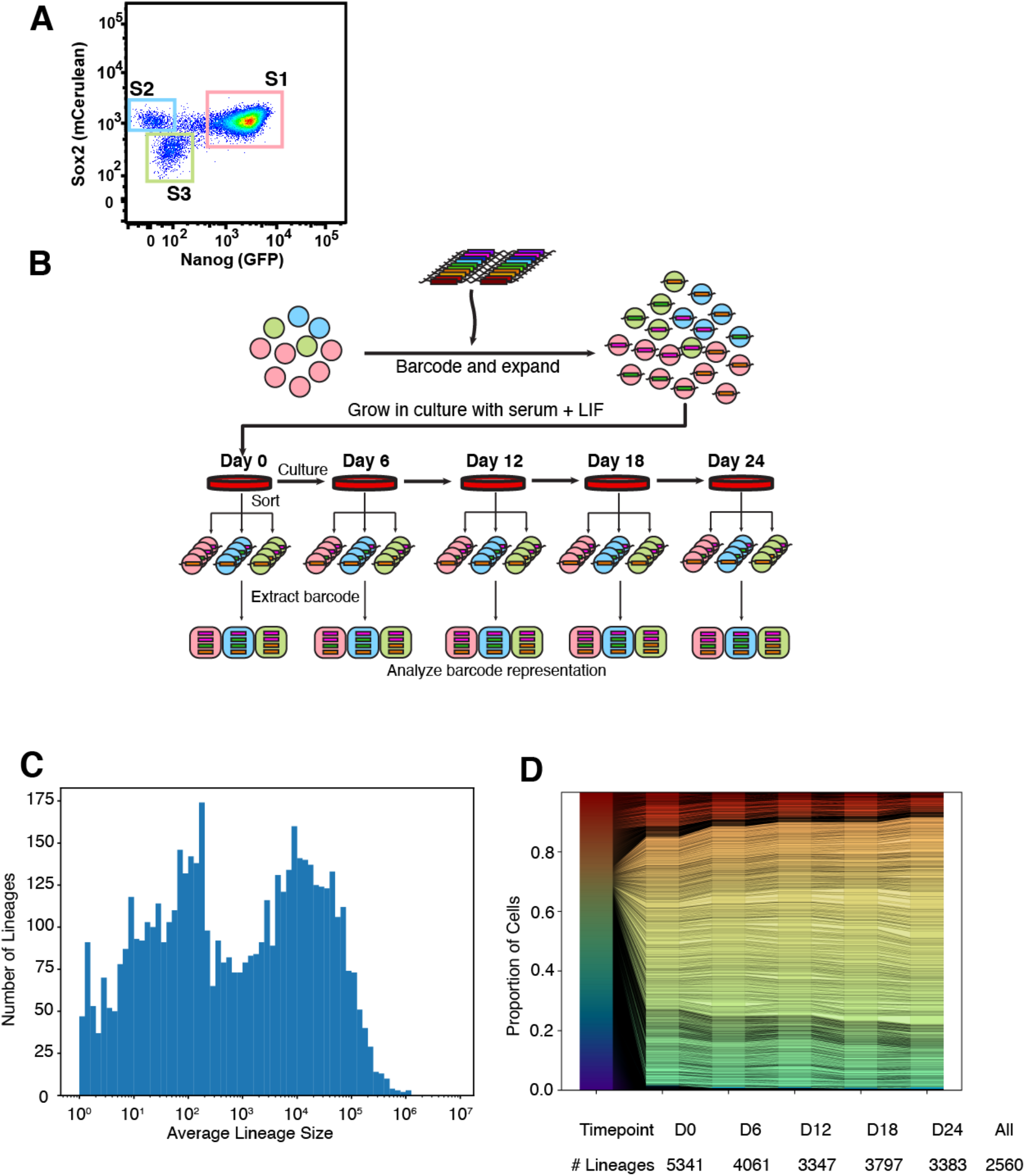
Generation and tracking of ESC lineages. **(A)** FACS plot showing the expression of Nanog and Sox2 in a population of mouse embryonic stem cells. Cells are binned into three states of expression (State 1 = Nanog High Sox2 High, State 2 = Nanog Low Sox2 High, State 3 = Nanog Low Sox2 Low). **(B)** Experimental Schematic. Lentivirally encoded barcodes are introduced into ESCs in the three States which are expanded into lineages. Cells are cultured over time, during which some ESCs switch between States. At the indicated timepoints half the culture is sorted into States 1-3 and the representation of each barcode (lineage) assessed in each state through sequencing (see Methods). Pink circles, Blue circles and Green circles represent State 1, State 2 and State 3, respectively. **(C)** Histogram showing the distribution of lineage sizes (in number of cells) on average across all timepoints. **(D)** Stacked bar plot representation showing the proportion each lineage contributes to the overall population across all timepoints. Each unique color row represents a distinct lineage. The number of lineages observed above background at each timepoint is indicated; 2,560 lineages are detected in at least one state at all timepoints. See also Figure S1.

We cultured these ESC lineages together and assessed their distribution across the three ESC states over a period of 24 days (Fig. 1B). First, we split our culture of 10^8^ cells, maintaining half in culture and isolating State 1, State 2, and State 3 cells from the other half by flow cytometric sorting for the fluorophore markers of Nanog & Sox2 (Fig. S1B). The cultured population was maintained at >= 2 * 10^7^ cells at all times to ensure high representation of lineages, and was split either every day or every other day due to the rapidly dividing nature of ESCs under standard culture conditions (doubling time ~12-14 hours). We assessed the distribution of ESC lineages across the states in a similar manner by sorting 10^8^ cells every 6 days (Figure 1B), isolating ~5*10^5^ cells for each state at each timepoint to ensure representation. We then identified the number of cells for each lineage in each state through the relative proportion of each barcode in each sample (Fig. S1C and Methods). We confirmed lineages were adequately detected through subsampling the data and noting a minimal effect on the size distribution of lineages detected (Fig. S1D). The estimated number of cells in each ESC lineage is shown (Fig. 1C, Fig. S1C, Methods). 2,560 lineages were confidently identified in at least one state at all timepoints of the experiment and are the focus of subsequent analysis.

Interestingly, the distribution of lineage sizes did not change appreciably over time (Fig. S1) nor did any particular lineages come to dominate the mixed culture by size (Fig. 1D). This is in contrast to lineage competition in other biological systems such as cellular reprogramming or differentiation, in which particular clones dominate the population (Chan et al., 2019; Shakiba et al., 2019). Our system reliably allowed us to track thousands of ESC lineages and their distribution across states over an extended period of time.

### Distribution and transitions of ESC lineages over time

We assessed the distribution of each ESC lineage across states by calculating the fraction of cells in each state. We represent this data on a ternary plot, in which each dot is a lineage and its position indicates the relative composition of State 1, State 2, and State 3 in that lineage at each timepoint (Fig. 2A, Figs. S2A-B). For example, a lineage on the top right corner of this plot indicates that all cells were in State 1 and the lineage was not detected in the State 2 or 3 samples, and analogously a lineage on the bottom left or top left corner indicates a lineage only present in State 3 or State 2, respectively. As expected based on flow cytometry data (Fig. 1A), the majority of cells in each lineage were in State 1 (Fig. 2A). Additionally, many lineages (955 of 2,560) were detected in all three states across all timepoints (Fig. S2A). At all timepoints, a subset of lineages was detected as present only in one or two states, as evidenced by the continued presence of lineages at or near the edges of the ternary plot.

**Figure 2.**
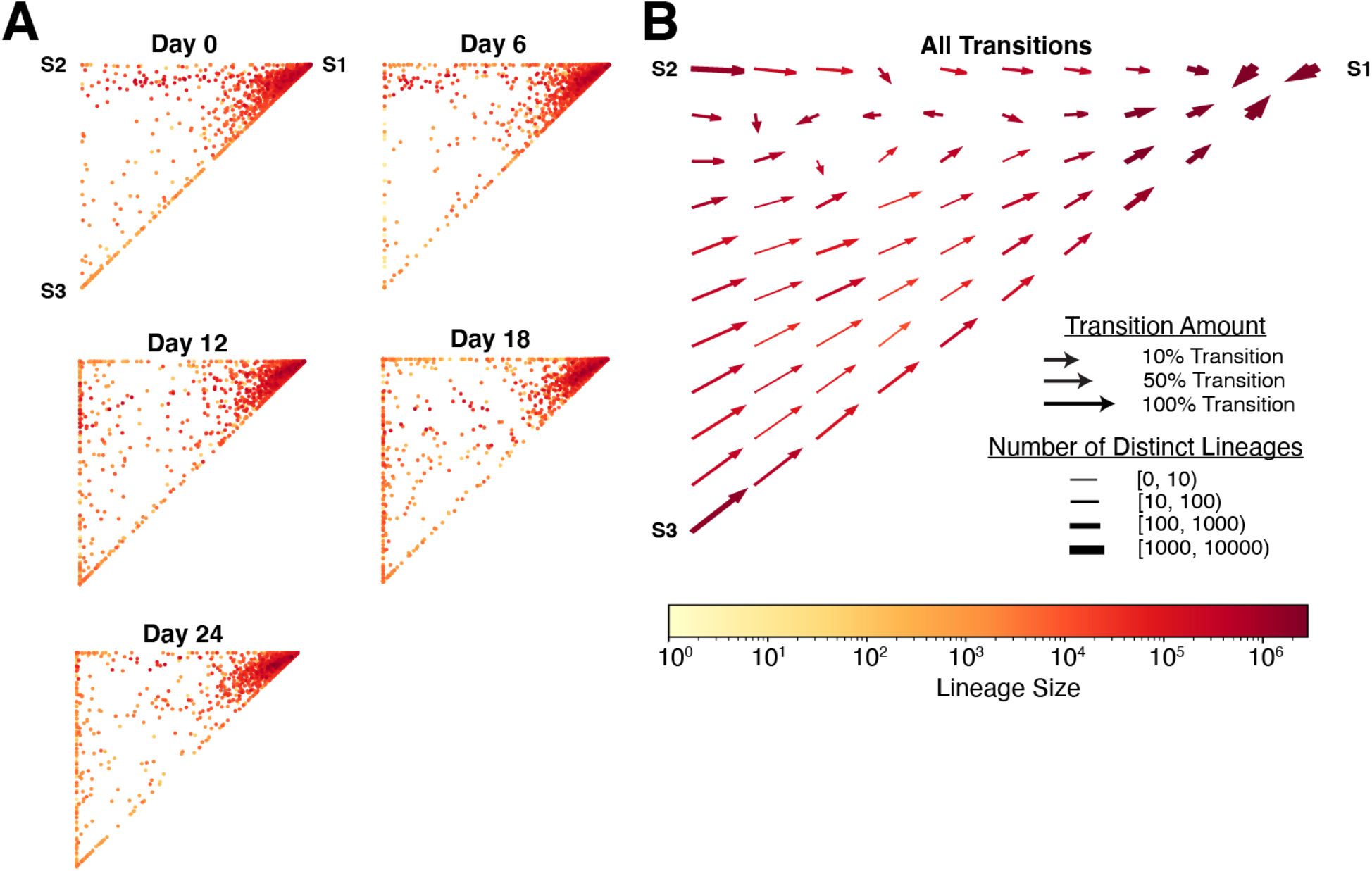
Distribution of ESC lineages over time. **(A)** Ternary plots showing the proportion of each lineage across states for all ESC lineages over time. Lineages in the corners are detected as present only within that one state. **(B)** Vector field showing the local average change in state proportions between two contiguous timepoints (such as Day 0 -> Day 6, representing a possible state transition). The local averages for all four transitions (Day 0 -> 6, Day 6 -> 12, Day 12 -> 18, Day 18 -> 24) are equally weighted. See also Figure S2.

Next, we sought to understand the dynamics of how cells in each lineage transitioned between states over time. First, we considered the change in proportion of each state for each lineage as a vector between two points on the ternary plot, and generated a vector field diagram. The diagram represents the summated transitions of all lineages present in each location of the plot: the fraction of lineages transitioning, the total size of cells, and the number of distinct lineages making transitions are all displayed (Fig. 2B). This plot was fairly constant for all four transitions captured in our experiment (Day 0 -> 6, Day 6-> 12, Day 12->18, and Day 18-> 24, Fig. S2C). The vector field plot revealed the overwhelming tendency of lineages present in the State 2 or 3 corner of the ternary plot to return to State 1 at the next timepoint, and for many lineages located in the State 1 region of the plot to switch into States 2 and 3. This is in agreement with previous studies showing individual ESC transitioning between Nanog-high and Nanog-low states (Filipczyk et al., 2015; Singer et al., 2014), albeit on a different timescale. However, the observation that ESC lineages also show net transitions between State 1 and States 2 & 3 at later timepoints is surprising, as cells transitioning in and out of a particular state might be expected to cancel out, leaving the lineage as a whole with no net change in position.

The information of how each individual ESC lineage transitioned between timepoints allowed us to calculate a matrix encompassing the transitional probabilities in this system. For each transition, we considered the proportion of each lineage in each state at time t_n−1_ as 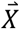 and those in each state at time t_n_ as 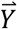. This allowed us to solve for the transition matrix **M** given by:

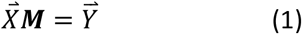

using least squares estimation (see Note). Transitional probabilities between the three states are shown (Fig. 3A), and the transitional probability matrix is given (Note). Next, we used this transition matrix (**M**) along with the known real sizes of each lineage (Fig. S1) before and after transition to calculate the net change in cell number for each type of transition (e.g., State 2 -> State 1, State 1 -> State 3, State 2 -> State 2, etc.) on average across all lineages. This rate of change represents a net combination of growth, birth, and death events for cells making each type of transition or staying within their state (Fig. S3A, see also matrix **G** in Note). Interestingly, cells transitioning from State 1 to State 3 show a net growth-birth-date rate of 21%, meaning cells making this transition show much smaller apparent population sizes after transition. In contrast, cells making the reciprocal State 3 to State 1 transition show a net growth-birth-death rate of 847%, indicating they greatly increased in cell number. Together, the rates of state transition and growth-birth-death between States 1-3 constituted a description of the dynamics in this system on average across ESC lineages.

**Figure 3.**
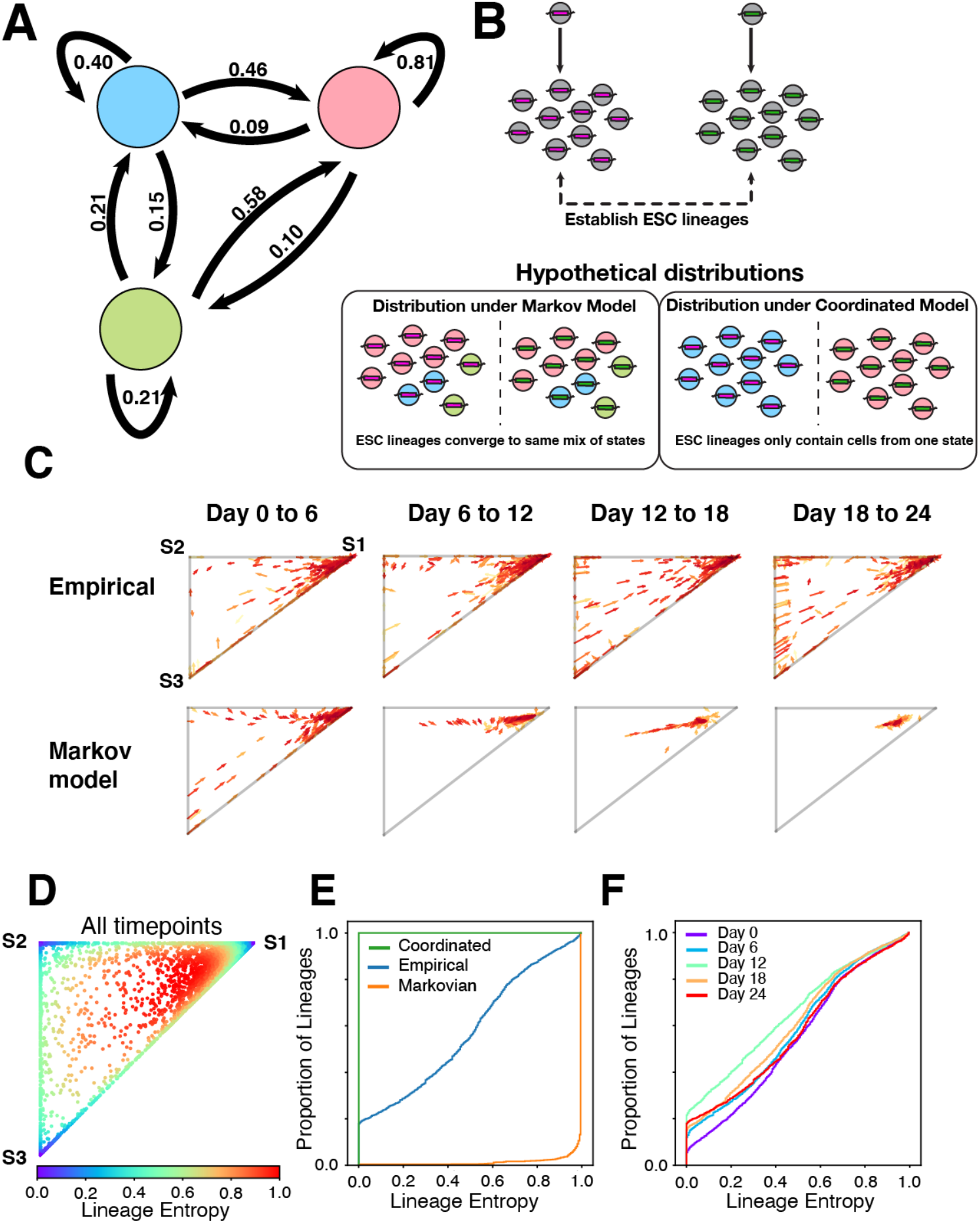
Some ESC lineages exhibit coordinated transitions between states in a manner that conserves entropy distribution. **(A)** Overall transitional probabilities between states inferred in the ESC system (see Note). Pink circle, Blue circle and Green circle represents State 1, State 2 and State 3, respectively. **(B)** Schematic of state transitions under two contrasting models. In the first, ESCs are assumed to possess the Markov property and are agnostic to their history, therefore over time ESC lineages converge to the same mix amongst states and same point on the ternary plot. In the second, ESC transitions are determined by their history and kin relations, therefore ESC lineages exhibit coordinated transitions and converge to the corners of the ternary plot. **(C)** Arrow plots show the transition of a subset of lineages from both the empirical data and that predicted by the Markov model. Arrow color indicates lineage size (scale matches that used in Figure 2). **(D)** Ternary plot displaying the lineage entropy (informational entropy) value for all lineages from all timepoints. **(E)** Cumulative distribution function (CDF) plot showing lineage entropy distribution for Coordinated and Markovian hypothetical models (green and yellow respectively) and the empirical data from D (blue). **(F)** CDF plot showing the distribution across all ESC lineages of lineage entropy for the empirical data at different timepoints.

### Path dependence and violation of a memoryless (Markovian) assumption

We sought to assess whether ESC lineage transitions had the Markov property. We consider a lineage in a given state (State 1, 2, or 3) if a plurality of cells in that lineage occupy the state; in other words, for each lineage whichever state contains the highest proportion of cells is defined as the state of that lineage. A stochastic process is said to possess the Markov property if for the set of variables under consideration ***X*** (in our case, the group of ESC lineages) occupying states given by ***S*** (States 1,2,3):

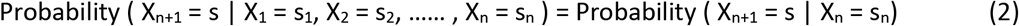

where sn are the states of each lineage at timepoint n (days 0, 6, 12, 18, and 24). Stated, this means that the distribution of ***X*** (lineages across states) at the next timepoint depends only on the present state, and not the entire history of transitions (see Note). In other words, the Markov property means where a lineage transitions next depends only on where it is now and not where it has been previously. We enumerated the probability of transitions between states for all 2,560 ESC lineages (Supplemental Table 1) to assess this statement. Strikingly, many lineages showed highly divergent conditional probabilities when the entirety of their history was taken into account (Supplemental Table 1). For example, we compared two lineage histories that were both in State 2 on Day 18 and assessed their probability of remaining in State 2 on Day 24. In the first history, lineages that were in State 1 on Days 0, 6, and 12 showed only a 21 percent probability of remaining in State 2 on Day_18->24_. However, this probability rose to 64 percent in the second history, where lineages were in State 2 on Days 0, 6, and 12 (p-value 1.47*10^−6^, Fisher’s exact test). The pathway of transitions for many lineages was highly proscribed, with cells following together through several transitions (Fig. S3B, Supplemental Table 1). Altogether, transitioning lineages were heavily biased to transition between States 1 and 2 or between States 1 and 3, with relatively few mixing transitions between States 2 and 3. This is consistent with the idea that States 2 and 3 represent distinct gene expression programs related to developmental timepoints downstream of State 1 (Chakraborty et al., 2020).

Further, we inferred transitions at the level of individual cells. If ESC state transitions possess the Markov property at the level of individual cells, distinct lineages of related ESCs should converge to the same distribution across states as every cell makes a separate choice of state regardless of its history and, therefore, its kin relations. This is equivalent to the idea that in a Markov process, each cell will sample from the same underlying probability distribution when choosing its next state. Thus, we compared ESC lineages over time under a Markov model with a coordinated model, in which related cells remain more likely to occupy similar states at later timepoints (Fig. 3B). We visualized the dynamics of lineages transitioning under both models (Supplemental Movies 1 and 2). A Markov model did not capture the dynamics of lineages in transitioning between states, as at all timepoints some lineages were distributed away from the equilibrium point and others appeared to be transitioning away (Fig. 3C, see also Fig. 2A, Fig. S2A). Additionally, a fully coordinated model did not capture the data as not all lineages were distributed on the corners of the plot, with most lineages containing cells in each state. Instead, the system appeared to contain a mix of ESC lineages retaining information about their kinship history and transitioning together and other lineages that were either not transitioning between states or had relatively equal numbers of cells making reciprocal transitions (i.e. one cell of the lineage transitions State 2 -> 1 while another cell transitions State 1 -> 2 such that the net proportion of the lineage in each state remains unchanged). Our system did not allow us to distinguish between these two possibilities. Nevertheless, together with assessment of state transition probabilities this analysis demonstrated ESCs do not transition between states in a completely memoryless manner. Rather, at least a subset of ESCs retain information about past states that influences future transitions.

### Informational Entropy in ESC lineages

Measuring how different lineages of cells distribute across state space represents a measure of information contained in the system. We sought to quantify the information retained by the system of ESC lineages. Compared to an equilibrium where all lineages become perfectly mixed in their proportion of states over time, the informational gain can be thought of as the relative information, information entropy, or relative entropy; we will use the term lineage entropy in the present context. All of these terms represent a quantity that approximates how far the system is from maximal uncertainty, which is achieved when all lineages are at a perfectly mixed equilibrium point. In the scenario of a Markov process, convergence of all related cells in an ESC lineage to an equilibrium distribution across states represents maximal lineage entropy. Conversely, ESC lineages where all cells are in the same state would represent minimal lineage entropy. We quantified the relative entropy of each lineage compared to the equilibrium point using a modified version of the Kullback-Leibler divergence ((Baez and Pollard, 2016), see Note). The lineage entropy at each point in the ternary plot is shown (Fig. 3D).

Next, we compared lineage entropy in our empirical data with that of the Markov model and a coordinated model, representing the data by plotting the cumulative distribution of lineage entropy across all lineages (Fig. 3E). We found the empirical distribution of lineage entropy diverged significantly from either model. More interestingly, the distribution of lineage entropy appeared relatively conserved over the timecourse of the experiment (Fig. 3F). Conservation of the distribution of lineage entropy is unexpected as the informational entropy in a stochastic system labeled at one distinct timepoint (introduction of barcodes) would be expected to strictly increase as the labels become diluted over time due to cells switching states independent of lineage history. This raises the question of whether this system is best considered stochastic or may be governed by additional variables.

### Defining ESC state by transition probability instead of gene expression

In examining the transitions of ESC lineages between states, we noted that most lineages fell into one of two categories: either they exhibited concerted transitions between State 1 and States 2 & 3 (Supplemental movie 1, red lineages) or they exhibited no net transitions at all (Supplemental movie 1, blue lineages). This led us to consider whether lineages might be properly classified on the basis of whether they were highly dynamic and exhibited concerted transitions between states (high “motility”) rather than their levels of gene expression (*Nanog* and *Sox2*). We calculated the motility of each lineage as the total distance travelled on the ternary plot over time (see Methods). First, we addressed whether the same lineages were transitioning into and out of State 1 over time. A subset of highly dynamic lineages displayed high motility in transitioning between states (Fig. 4A, Fig. S5A). These lineages did not necessarily represent the smallest sized lineages and in fact showed an almost normal distribution with respect to lineage size (Fig. S5B). Analyzing motility of all lineages over time demonstrated a consistent subset transitioning between states, resulting in a ‘sawtooth’ type appearance when motility was plotted against time (Fig. 4B, Fig. 4C, and Fig. S6A). We confirmed highly dynamic lineages were not due to sampling a portion of cells in states by considering the change in motility as data was sub-sampled (Fig. S6B), which was minimal. To visualize this effect in another fashion, we plotted the portion of each lineage transitioning at each timepoint and colored lineages by their overall transition amount across the whole experiment (Fig. 4D, Fig. S6C). The consistent ‘skew’ of red, highly dynamic lineages to the right indicates that this same group of lineages was transitioning between states at each timepoint. Finally, we calculated the pairwise correlation of motility for each lineage across transitions, which confirmed a relationship between motility at early transitions and later transitions (Fig. S7A).

**Figure 4.**
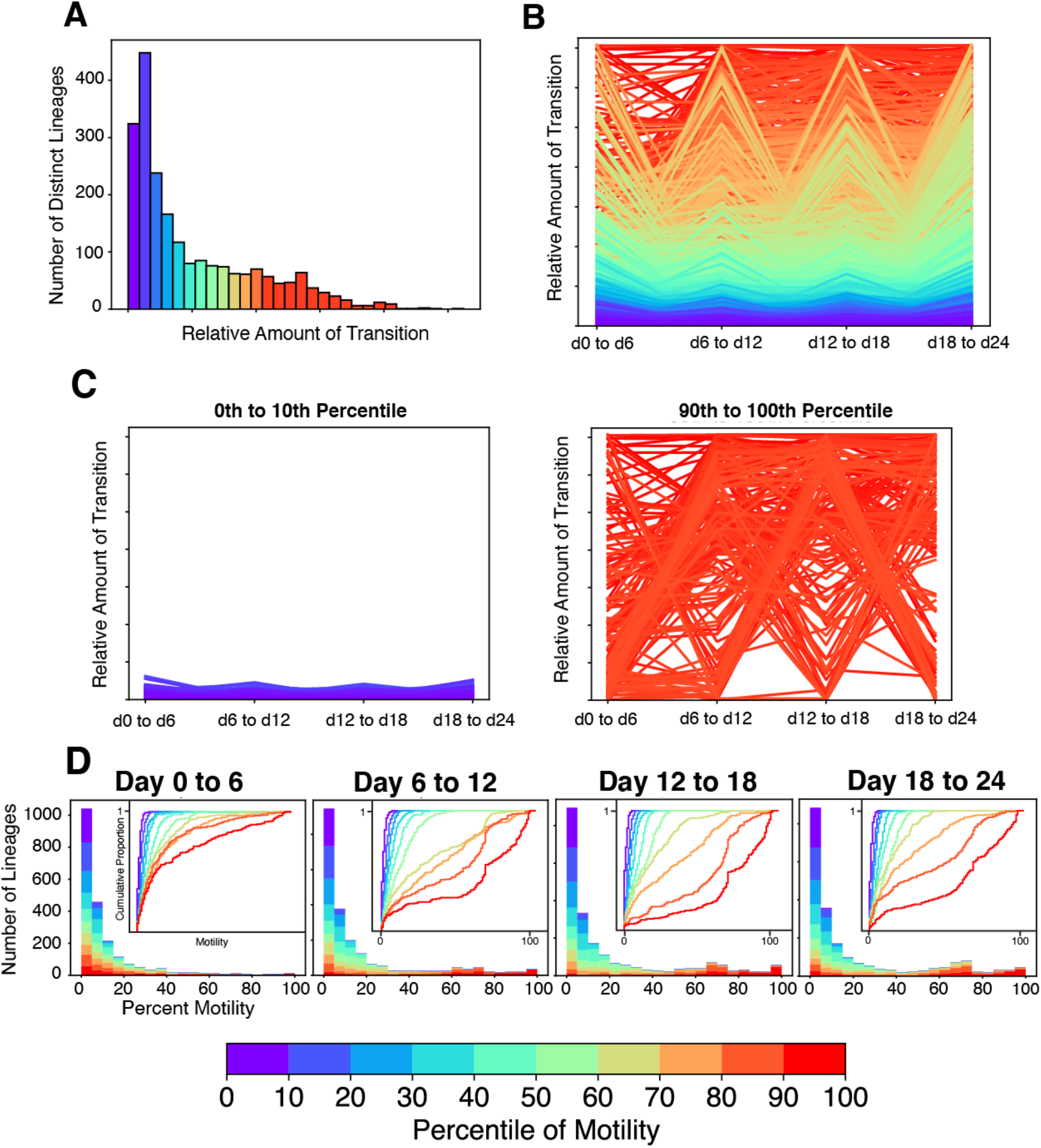
A consistent subset of ESC lineages is characterized by a high amount of state transitions. **(A)** Histogram showing the number of lineages with differing amounts of transition between states. Lineages are colored according to their percentile rank of their motility between states across all timepoints relative to all 2,560 measured. **(B)** Line chart showing the amount of transition for all lineages at each transition point. **(C)** Line charts showing the amount of transition for lineages in the first decile (left) and the last decile (right) of motility. **(D)** Stacked bar plots show the distribution of percent motility from lineages in different decile groups of transition. Inset: CDF plot providing additional visualization of distribution of percent motility. In A-D, all lineages are decile ranked according to their cumulative overall motility across transitions and colored identically in each panel.

This led us to consider whether motility was a conserved feature defining state in the ESC system, and if states would be better considered as ‘motile’ lineages and ‘non-motile’ lineages irrespective of specific Nanog and Sox2 levels. When dividing lineages into motile and non-motile states, we again did not observe the Markovian property, as high motility at all prior transitions was associated with a higher probability of remaining high in motility when compared against a lineage that was only high in motility at the immediately preceding transition (Supplemental Table 2, Fig. S8, and Note). To further elucidate whether motility of a lineage was a conserved feature, we performed a repeat of our entire experiment in duplicate (Fig. S7B) and compared the motility of each lineage at each transition between replicate experiments, comparing lineages with the same barcode in each replicate to each other. We found a small correlation (Pearson R = 0.22) between motility across replicates (Fig. S7C), which was significant when compared to a model in which each lineage sampled its motility randomly from the experimental distribution (Fig. S7D and Note) but less than the average correlation of motility within each replicate (Pearson R ranging from 0.18 to 0.32). Additionally, both replicates demonstrated non-Markovian state transitions and conservation of lineage entropy (data not shown). Together, we conclude that motility between cell states shows modest correlation across transitions and replicates in the ESC system.

## Discussion

We analyze lineages of ESCs transitioning between states over time and find a subset of lineages with cells that transition between states together. These lineages appear to follow distinct, proscribed paths of state transitions, with the full history of the lineage influencing the probability of future transitions. Therefore, we deduce at least some ESC lineages do not possess the Markov property of memorylessness. Interestingly, the number of such lineages and the degree to which they differ from a randomized Markovian model of transitions appears relatively conserved over time.

Our results contrast to some extent with previous reports indicating that ESC state transitions may occur in a manner independent of lineage history (Filipczyk et al., 2015; Singer et al., 2014). The previous reports measured ESC transitions over shorter timescales (hours) than the 24 days measured here, which may in part account for this difference. Fluctuations between states in biological systems has previously been proposed to arise in part from slow global fluctuation of the transcriptome, possibly over timescales as long as a week (Huang, 2009). The results presented here would support such a model. While the source of such fluctuations is unknown, one possible source could be oscillators that may in part drive state transitions, and identification of such systems will be of great interest.

The idea that kin related cells transition between states in a correlated fashion suggests the presence of as yet unknown hidden variables that may govern these transitions. We defined state in this study on the basis of Nanog and Sox2 gene expression, and while our previous observations suggest this captures the greatest component of heterogeneity in ESCs (Chakraborty et al., 2020), this is still a relatively limited two dimensional reduction of gene expression space. Perhaps defining the full transcriptome of ESCs along with lineage information will allow better prediction of future states. On the other hand, a recent study of hematopoiesis *in vitro* and *in vivo* captured both full transcriptomic information and lineage for single cells, yet found sister progenitors to have intrinsic fate biases that could not be accounted for by the transcriptome (Weinreb et al., 2020). Many elegant systems of encoding kinship relations exist in biology, and understanding how ESCs may use such systems and to identify kin alongside developing a more complete picture of how cells encode their histories will be of great interest.

The results here may also have implications for cellular competition. Development in the mammalian body is known to occur in part through competition, whereby the fittest precursor cells survive and make greater contributions to the adult organism (Claveria et al., 2013; Dejosez et al., 2013). Greater fitness in cell competition experiments has often been attributed to the ability of elite clonal lineages to rapidly divide, thereby increasing their number (Shakiba et al., 2019). We find a subset of ESC lineages shows consistently high motility in transitioning between states but does not appreciably increase its growth rate and thus does not dominate the population. This may indicate that in some contexts highly dynamic lineages are those with a greater ability to switch between states in a manner distinct from elite growth ability.

Many biological systems show a constant homeostatic ratio between cell types or cell states. For example, intestinal crypts or epidermis in the skin maintain a constant proportion between stem cells at the base of the tissue and more differentiated cells towards the periphery. Informational entropy can be considered in many ways when examining such a system. In the present study, we examine informational entropy at the lineage level, analogous to study of how particular stem cells and their descendants partition across states over time. We find a consistent distribution of relative lineage entropy in ESC state conversions, meaning though some ESC lineages skew into particular states and others mix across states, the ratio between these types is preserved. The analogy in a tissue would be if not only particular stem cells were more stem-like than other stem cells more primed towards differentiation, but also the system overall preserved the ratio between these types of stem cell lineages. Identifying whether this property holds in biological development and homeostasis or is limited to the system of cultured embryonic stem cells studied here will be of interest. Together, the results presented here showing ESC state transitions are non-Markovian for some lineages may have implications for understanding the flow of biological information during state transitions and may suggest homeostasis and biological robustness are best understood both at the level of individual cells and at the level of the lineages from which they are descended.

## Supporting information

Methods and Theory Notes

## Acknowledgements

This work was supported in part by the Koch Institute Support (core) Grant P30-CA14051 from the National Cancer Institute. We thank the Koch Institute’s Robert A. Swanson (1969) Biotechnology Center for technical support, specifically Flow Cytometry and the MIT BioMicro Center. SH acknowledges support from NIH T32 GM007287-45. SG acknowledges support from NIH NCI K08 CA237856 and a Charles W. and Jennifer C. Johnson Clinical Investigator Award (MIT).

## Author contributions

SG designed the study. TU performed experiments, generated the models, and interpreted data. SH generated the cell lines and interpreted data. SG wrote the manuscript with input from TU and SH.

---

**Figure S1.**
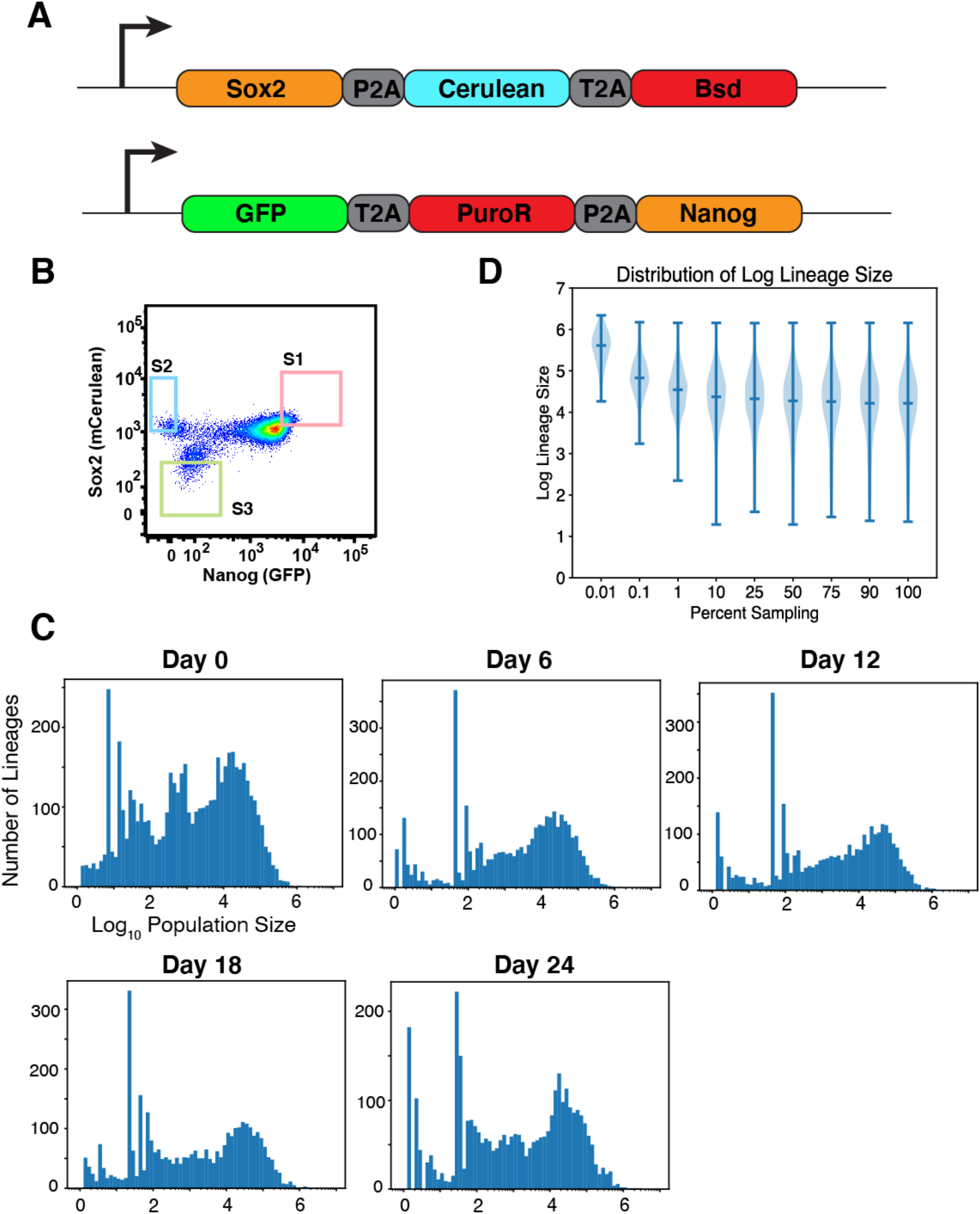
ESC fluorophore targeting and lineage sizes. **(A)** Schematic of loci for generation of *Sox2-P2A-Cerulean* and *GFP-P2A-Nanog*. *BSD* and *PuroR* refer to genes encoding resistance to the selectable markers blasticidin and puromycin, respectively. Insertions were verified heterozygous by locus PCR. **(B)** Schematic showing representative gates used to sort populations for States 1-3 at each timepoint of the experiment. Sorted cells were analyzed for >99% purity from the other two gates. **(C)** Distribution of ESC lineage sizes at each timepoint of the experiment **(D)** Distribution of lineage sizes as data (reads) are sub-sampled. Percent sampling refers to the proportion of randomly selected reads from the fastq file that were retained for analysis, and the distribution across inferred lineage sizes is shown. The median (center line) and 1^st^-99^th^ percentiles (whiskers) are indicated.

**Figure S2.**
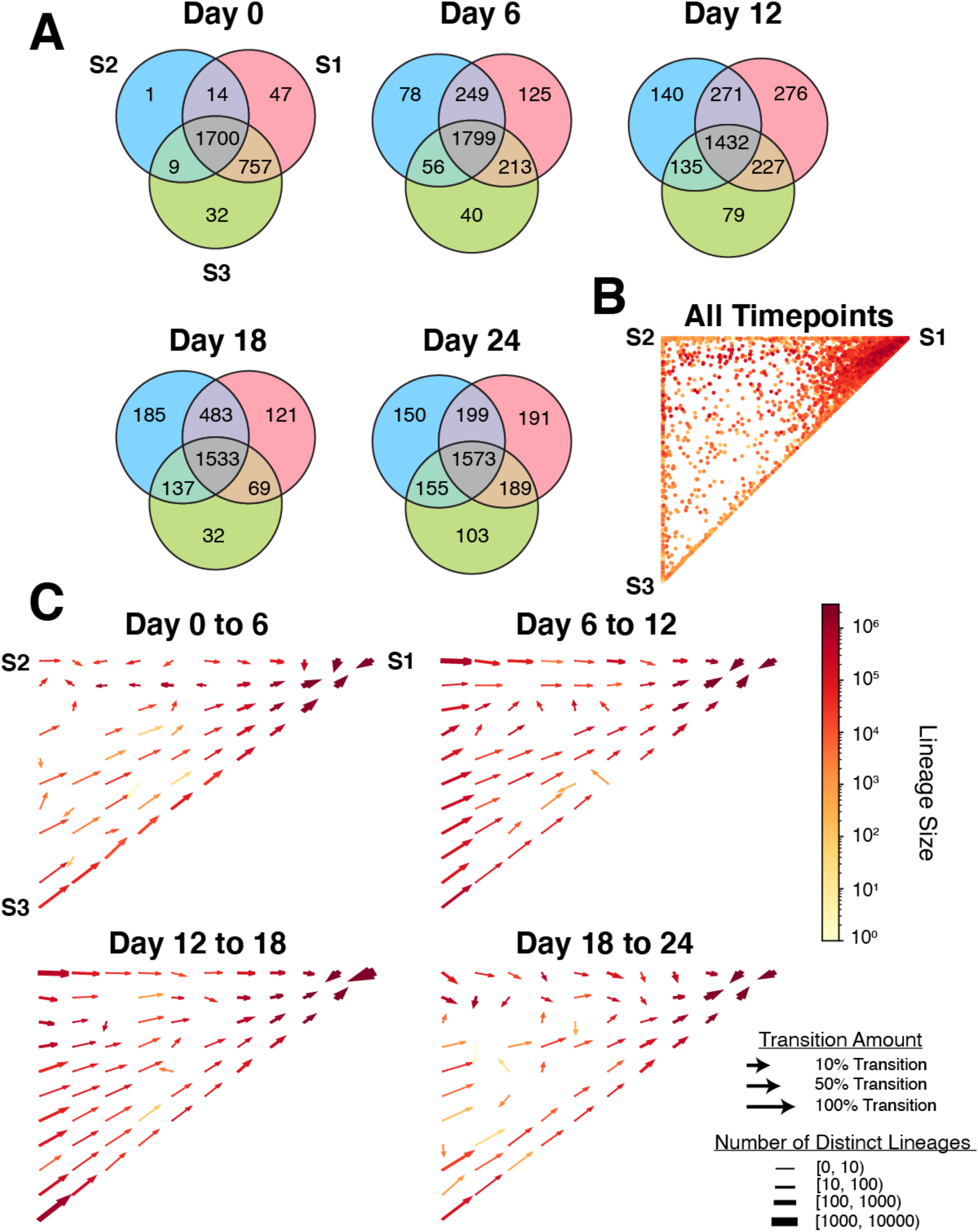
Number and transitions of lineages between states. **(A)** Venn-diagrams showing number of lineages that are detected in each state or in multiple states at all timepoints. **(B)** Ternary dot plot showing the distribution of lineages across three states for all lineages at all timepoints combined. **(C)** Vector field plots showing the local average change in state proportions (state transitions) for each of the four measured transitions between timepoints in the experiment. Color bar for panels B,C indicate lineage size in number of cells.

**Figure S3.**
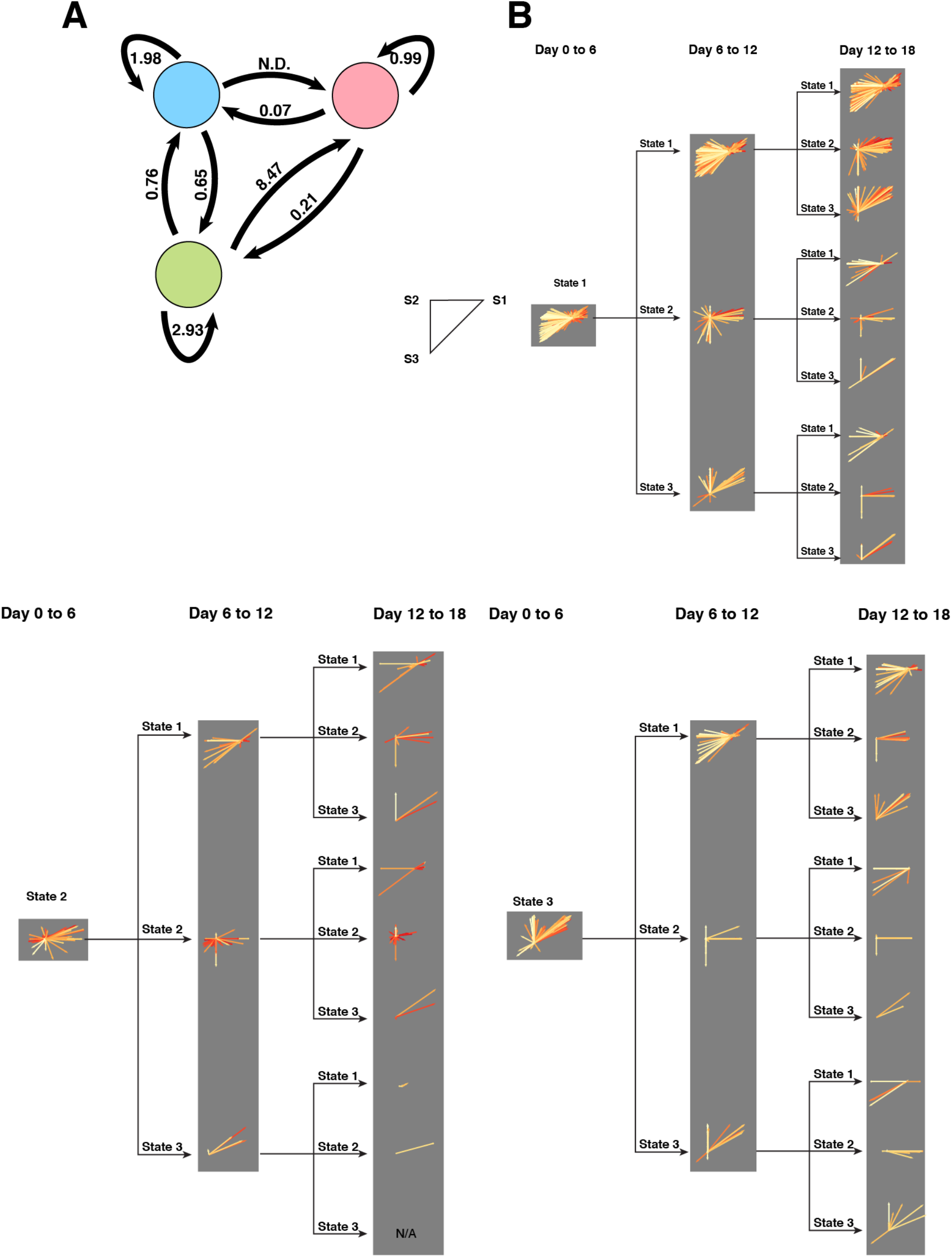
Average growth-birth-death rate of ESC and transitions of discrete lineages in different states. **(A)** State diagram showing all possible cell state transitions and the growth-birth-death rate net of state transitions associated with each transition (see Note). For example, cells remaining in State 2 show a net growth-birth-death rate of 1.98, meaning such cells nearly double in size between timepoints. Pink circle, Blue circle and Green circle represent State 1, State 2 and State 3, respectively. **(B)** Recenter plots showing how lineages in different states transition over time from Day 0 to Day 18. Each lineage is assigned a state (S1,S2,S3) on Day 0 based on where a majority of its constituent cells are detected, and is represented by a vector. Each plot follows a collection of transition vectors from lineages in a particular state that follow a particular set of transitions across time. These vectors are then recentered at the origin point (0,0) of the group of vectors. Vector direction indicates the transition of each lineage with respect to a ternary plot (shown at left) across the indicated timepoints. For example, a lineage in S1 at day 0 represented here by a left arrow vector (−1,0) indicates this lineage transitions entirely to S2, and a lineage represented by vector (−0.5,−0.5) would indicate an S1 lineage transitioning equally to States 2 and 3. The length of each vector correlates with the change in lineage size, and coloration indicates the present size. Color scale matches that used in main Figures 2 & 3. Gray background provided for ease of visualization.

**Figure S4.**
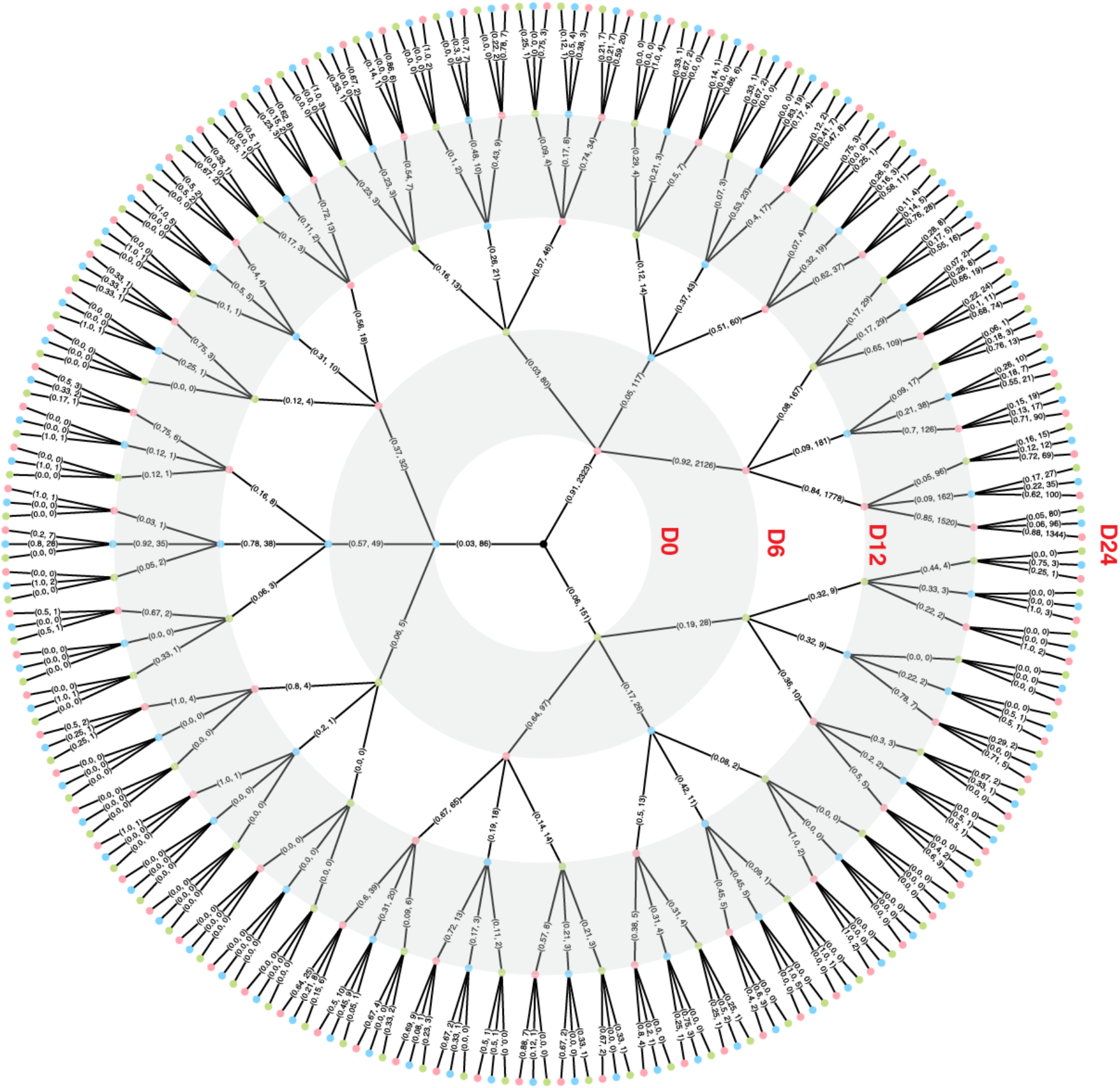
Probability decision tree of lineage Nanog-Sox2 state transitions. Diagram showing the likelihood of state transitions of ESC lineages in particular Nanog-Sox2 states at all timepoints. Day 0 is represented as the innermost dot, moving outward to Days 6, 12, 18, and 24. Pink, blue, and green dots represent States 1, 2, and 3, respectively. Numbers represent the conditional probability of transition and the number of lineages making said transition down that branch as a (x,y) pair.

**Figure S5.**
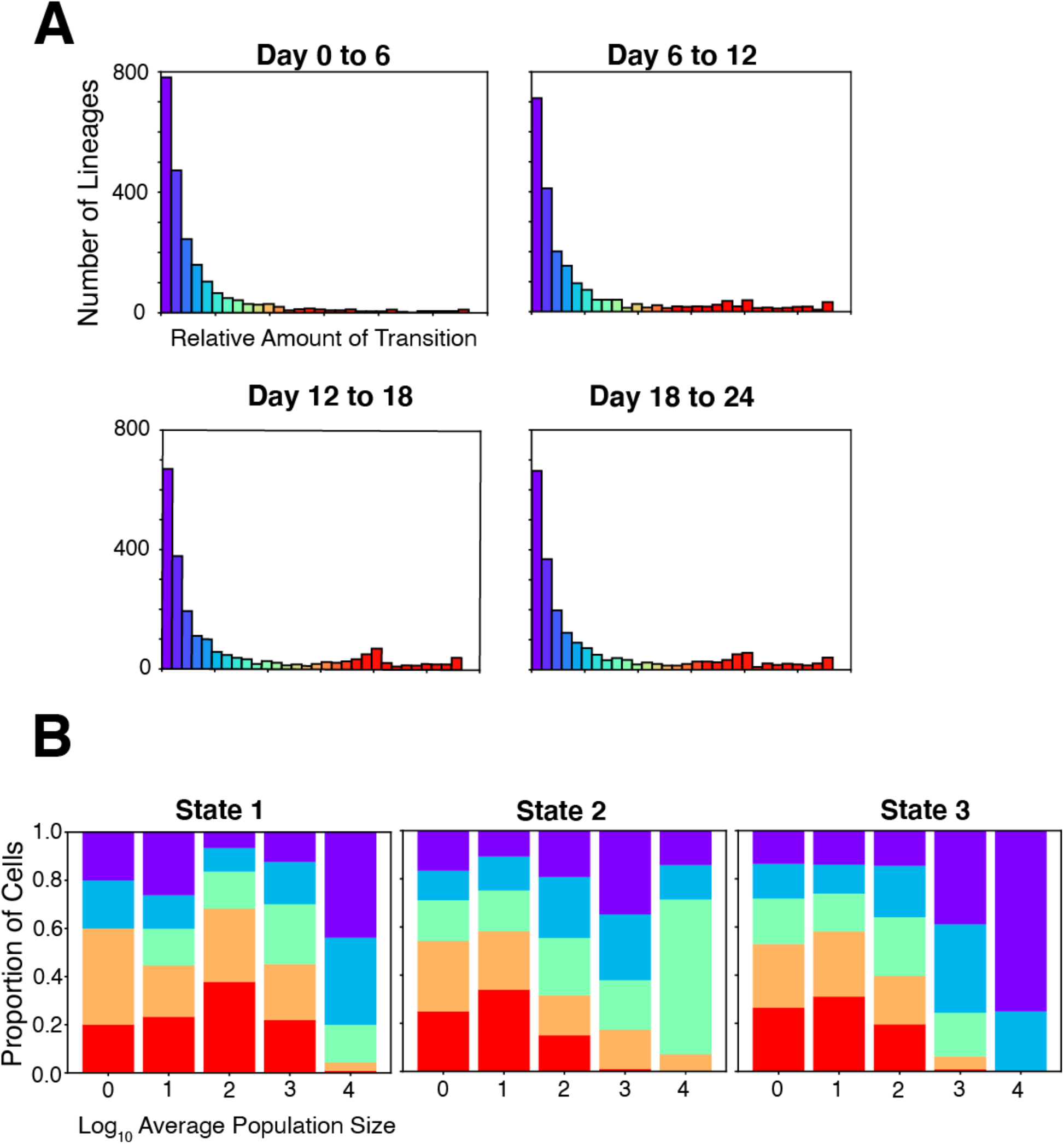
Motility analysis of ESC lineages. **(A)** Histograms showing the distribution of lineages with different amount of transition between states (motility) at each transition point. Lineages are colored as in main Figure 4. **(B)** stacked bar plots revealing the proportion of lineages of different sizes (binned by order of magnitude) with different motilities in each state, summed across all four transitions. The state of a lineage is defined as where a majority of its ESCs are located at the first timepoint of the transition. Lineages are grouped into 5 quintiles of transition: red (top 80-100^th^ percentiles), orange (60-80^th^ percentiles), teal (40^th^-60^th^ percentiles), blue (20^th^-40^th^ percentiles), violet (0^th^-20^th^ percentiles). Note the highest transitioning lineages (red) are not heavily skewed towards the smallest size lineages, arguing against the idea that a sampling effect alone could explain the observed data.

**Figure S6.**
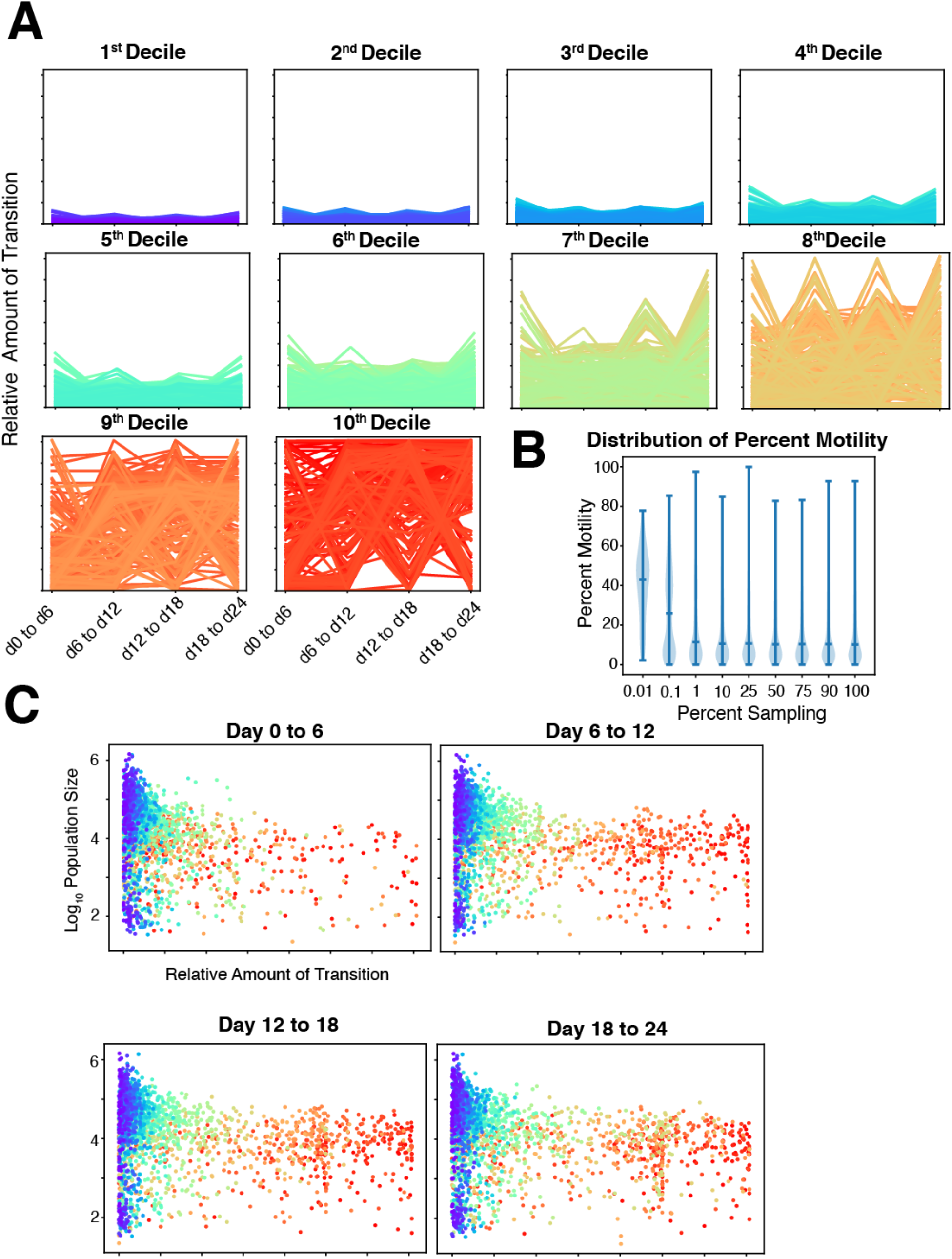
Decile motility plots and relationship between motility and lineage size. **(A)** Line charts showing the amount of transition for each lineage by decile of overall motility. 1^st^ and 10^th^ deciles are also shown in main Figure 4C. **(B)** Violin plot showing the distribution of lineage motility as data (reads) are sub-sampled. Percent sampling refers to the proportion of randomly selected reads from the fastq file that were retained for analysis, and the distribution of motility (percent transition) is shown. The median (center line) and 1^st^-99^th^ percentiles (whiskers) are indicated. **(C)** Scatter plots showing the amount of transition and sizes of each lineage in each transition. Each dot is colored by the overall amount of that lineage’s transition across all timepoints – colors match those used in Figure 4.

**Figure S7.**
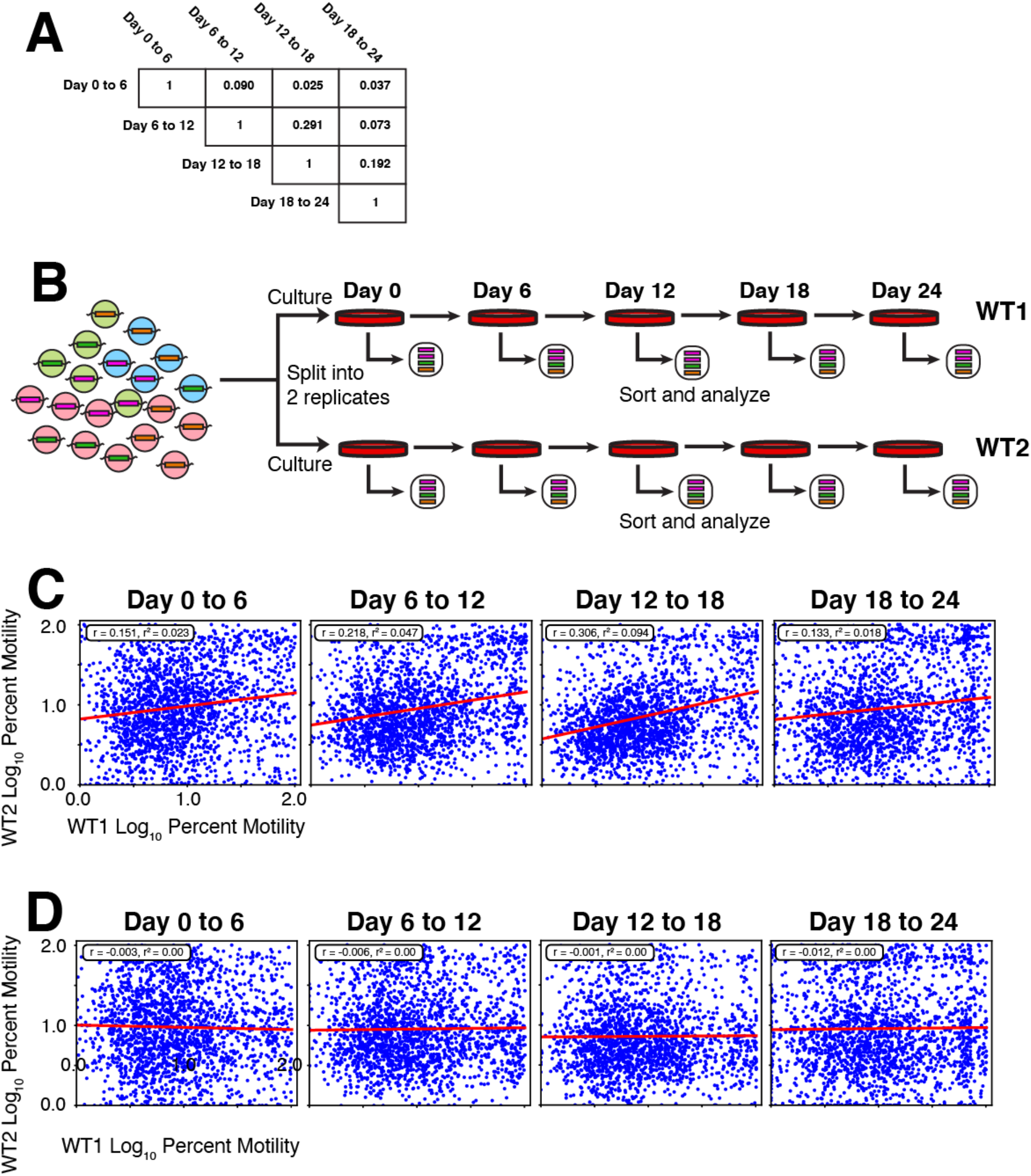
Correlation between replicates. **(A)** Matrix showing the Pearson correlation (r-value) for motility of all lineages when different transitions are plotted against one another. **(B)** Schematic of replicate experiment. The initial barcoded population of ESC lineages was split into two equal portions. Each was maintained separately over 24 days, and ESC lineages in States 1-3 were assessed by sorting representative pools from States 1-3 on Days 0, 6, 12, 18, and 24. Barcodes were sequenced and the relative proportion of each lineage in each state assessed. **(C)** Scatter plots depicting the relationship between percent transition amounts in replicate 1 (WT1) and replicate 2 (WT2) at all transitions in the experiment. The Pearson correlation (r-value) and r^2^ are shown. **(D)** Scatter plots depicting the relationship between percent transition amounts in randomized replicates. In each replicate, transition amounts were chosen randomly for each lineage from the empirical distribution of transition amounts and then plotted against each other. No significant correlation is observed.

**Figure S8.**
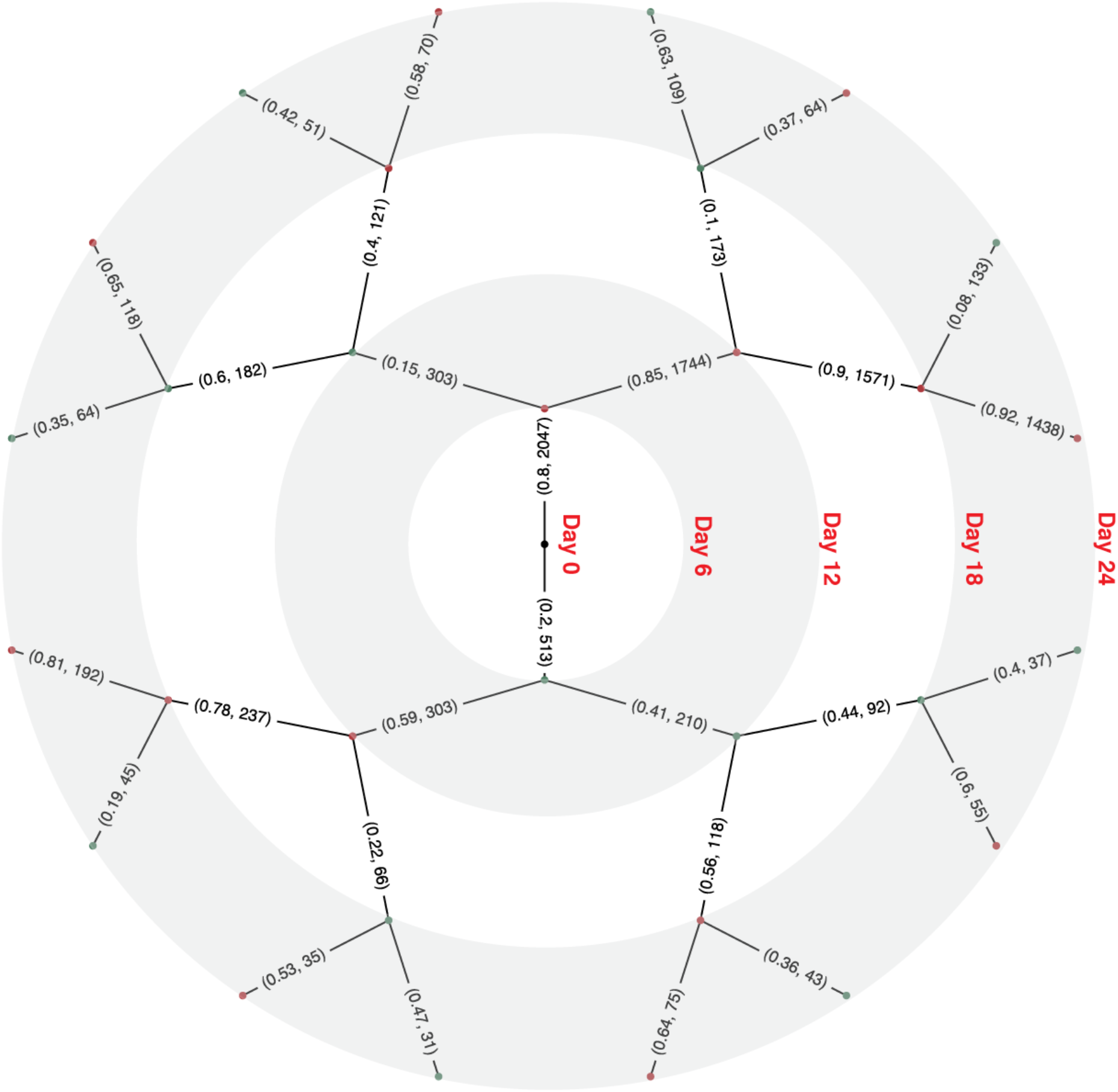
Probability decision tree of lineage motility state transitions. Diagram showing the likelihood of state transitions of ESC lineages in particular motility states at all timepoints. Day 0 is represented as the innermost dot, moving outward to Days 6, 12, and 18. Red and green dots represent lowly motile and highly motile states, respectively. Numbers represent the conditional probability of transition and the number of lineages making said transition down that branch as a (x,y) pair.

