## Supplementary material for "Lineages of embryonic stem cells show non-Markovian state transitions": Methods and Theory Notes

**Materials and Methods**

**Culture and labeling of embryonic stem cells**

**Cell line information**

Two mouse embryonic stem cell (ESC) lines were used in this study: V6.5 (gift from the Jaenisch Laboratory, Whitehead Institute, MIT) and V6.5 derived *Nanog-GFP/Sox2-mCerulean* generated as described below using stably-integrated DNA barcodes ((Bhang et al., 2015), Addgene #67267).

**Cell line maintenance**

Cells were cultured in liquid medium on 10-cm tissue plates pre-coated with 0.2% gelatin in phosphate-buffered saline (PBS). The liquid medium contains 415 mL of Dulbecco’s Modified Eagle Media (DMEM, Catalog number 1195073, Gibco), 5 mL of 0.1 mM L-glutamine, 5 mL of 0.1 μM non-essential amino acids, 5 mL of 0.1 μM penicillin-streptomycin antibiotics solution, 5 mL of 1 M HEPES buffer, 4 μL of 14.3 M beta-mercaptoethanol, 82.5 mL HyClone fetal bovine serum (FBS), and 55 μL of 1000U/mL Leukemia Inhibitory Factor (LIF). DMEM + additive components were filtered sterilized using a 0.45 micron filter (Catalog number 430770, Corning) before adding FBS and LIF.

**Fluorophore tagging of pluripotency genes**

Endogenous *Nanog* and *Sox2* genes were tagged by *GFP* and *mCerulean* via CRISPR-Cas9 induced homology directed repair (HDR). Single-guided RNAs targeting upstream of the start codon (*Nanog*) or downstream of the stop codon *(Sox2)*. The single-guide RNA sequence (For *Nanog* locus: 5’-CACCGTCAGTGTGATGGCGAGGGA-3’ and its complementary 3’-AAACTCCCTCGCCATCACACTGAC-5’; For *Sox2* locus: 5’-CACCGATTGGGAGGGGTGCAAAAAG-3’ and its complementary 3’-AAACCTTTTTGCACCCCTCCCAATC-5’) was cloned into PX330 plasmid using BbsI restriction site. The plasmid was then introduced to the cell using cationic lipid transfection (Lipofectamine 2000, Invitrogen, Catalog number #11668019) along with a homology-directed repair construct encoding the relevant fluorophore (*GFP* for *Nanog* and *mCerulean* for *Sox2*), T2A/P2A post-translational cleavage sequences, and a drug resistance gene (*Puromycin^R^* for *Nanog* and *Blasticidin^R^* for *Sox2*). Cells were then selected in culture medium with Puromycin and Blasticidin at concentrations of 2 μg/mL and 4 μg/mL, respectively, for 14 days.

**Molecular barcoding of ESC**

ESCs were labeled using ClonTracer library ((Bhang et al., 2015), Addgene #67267). *Nanog-GFP*; *Sox2-mCerulean* V6.5 ESCs were transduced with the aforementioned barcoding library by spinoculation. ~250,000 transduced cells were selected by the expression of RFP using flow cytometric sorting. Successfully transduced cells were then cultured for ~2 weeks until at least 20 million cells were present in the population*.* These cells were then cultured and sorted as described in Figure 1B.

**Sorting and Analysis**

**Flow cytometry and fluorescence activated cell sorting (FACS).**

Barcoded *Nanog-GFP*; *Sox2-mCerulean* V6.5 ESCs were analyzed for their Nanog-Sox2 expression on a BD LSRII HTS-2 with FACSDiva v8.0 acquisition software. Data gathered from flow cytometry were then analyzed by FlowJo V9.9. Live cells were first selected based on the forward scatter area (FSC) vs. side scatter area (SSC). Single cells were then selected based on forward scatter height (FSC-H) vs. forward scatter width (FSC-W). The expression of GFP and mCerulean (a proxy for Nanog and Sox2 expression, respectively) were observed using FITC and Pacific Blue detector channels using wild type V6.5 or singly GFP/mCerulean labeled ESC as compensation controls. For the experiment detecting ESC lineages, this barcoded fluorophore-tagged ESC line was sorted by a BD FACS ARIA machine into 3 states based on the level of GFP (representing Nanog level) and mCerulean (representing Sox2 level). Distinct states were sorted into culture medium, spun at 223 rcf for 5 minutes, and cell pellets frozen for analysis. FACS analyzers and sorters utilized were provided by The Swanson Biotechnology Center Flow Cytometry Facility, Koch Institute for Integrative Cancer Research at MIT, MIT.

**Extracting DNA barcodes from ESC genome**

Genomic contents from sorted cells were extracted using Sigma GenElute Mammalian Genomic DNA Prep Kit (Catalog number #G1N70) and PCR amplified using the primers listed in Supplemental Table 4. DNA samples from different states and different timepoints were amplified using unique reverse primers. PCR reactions contained 10 μL of 5X NEB Phusion High-Fidelity buffer (Catalog number B0518S), 1.5 μL of DMSO, 1 μL of dNTPs (NEB, catalog number N0447S), 1 μL of 10 μM forward primer, 1 μL of 10 μM reverse primer, 0.5 μL of NEB 2000U/mL Phusion High-Fidelity DNA Polymerase (Catalog number M0530S), genomic DNA, and ddH_2_0 to 30 μL. Reactions were amplified using an annealing temperature of 55.5ºC for 25 cycles. Amplicons were pooled together for DNA next-generation sequencing.

**Illumina sequencing**

Amplicons were sequenced using Illumina MiSeq and NextSeq500 sequencers. This sequencing service was provided by MIT BioMicro Center, MIT Department of Biology. The flowcell primer 5’-CCGAGATCTACACACTGACTGCAGTCTGAGTCTGACAG-3’ was used.

**Informatic read processing**

**Data and code availability**

The fastq files for this experiment are available on Sequence Read Archive (SRA) with Bioproject: PRJNA670562. Python scripts for processing Fastq files generated by sequencing and subsequent analyses are available on GitHub (https://github.com/teeu97/lineage-entropy).

**Quality metrics for sequencing reads**

Sequencing reads in Fastq files generated from DNA next-generation sequencing were quality filtered by Phred Score (Phred +33, Illumina 1.9) and for sequencing errors compared to the reference amplicon. The reference sequence is: 5’NNNNNNNNNNNNNNNNNNNNNNNNNNNNNNAGCAGAGCTACGCACTCTATGCTAGTGCTAGAGATCGGAAGAGCACACGTCTGAACTCCAGTCACXXXXXXXXXXATCTCGTATGCCGTCTTCTGCTTG-3’ where N represents barcodes and X sample indices, respectively. Before used in subsequent analyses, each read must pass the following criteria: 1. The sum of the Phred quality values of the barcoding region (the first 30 bases) must be greater than 80% of maximum Phred quality values (40 Phred score/base * 30 base = 1200 maximum Phred score); 2. The Hamming distance (the number of base mismatches) between the read and the reference sequence in the first constant region (the 31^st^ to 95^th^ nucleotide) must be less than 6; 3. The Hamming distance between the read and the reference sequence in the second constant region (the 106^th^ to 130^th^ nucleotide) must be less than 3; 4. The Hamming distance between the read’s sample index and one of the sample indices must be less than 2. The number of reads that passed these quality metrics are listed in Supplemental Table 3. Reads that passed the quality metrics were then separated based on sample indices. (Note that each sample was amplified by a unique reverse primer that contained a distinct sample index. This allowed us to separate them in this pipeline.) Finally, reads that passed the quality standard with barcodes less than 6 Hamming distance away from each other were collapsed together.

**Total count normalization method**

Quality filtered reads were then normalized into cell counts based on the State sample from which they originated. Specifically, reads that were in State 1 were normalized to 90,000,000 cells whereas State 2 and State 3 reads were normalized to 5,000,000 cells. This ratio of State 1: State 2: State 3 = 90:5:5 was approximated from the distribution of cells in the FACS plot (Figure 1A) and normalized to the total of 10^^8^ cells sorted. Please note that this calculation approach allows samples with different sequencing depth to normalize to the same total number of cells if they represent cells in the same State. For example, all State 1 cells from Day 0, 6, 12, 18, and 24 are always normalized to 90,000,000 cells. Lineages are further considered if and only if the sum of their read numbers in State 1, State 2, and State 3 is greater than 0 at all timepoints.

**Motility of ESC lineages**

Relative transition amount or motility is defined by the Cartesian distance between the location of a lineage in the first timepoint (x_1_, y_1_) to the location in the second timepoint (x_2_, y_2_) on the ternary plot. In other words,

**Motility** = $\sqrt{{(x_{1}-x_{2})}^{2}+{(y_{1}-y_{2})}^{2}}$

Using ternary plot coordinates for (x,y) where a lineage entirely in State 1 is (1,0), State 2 is (0,1), and State 3 is (0,0). Note that the motility is always between 0 (no change in state proportions between two timepoints) and $\sqrt{2}$ (100% change along State 1-State 3 axis). The maximum change of state proportions along State 1-State 2 axis and State 2-State 3 axis is 1. This weighting was regarded as appropriate given the greater gene expression differences between States 1 and 3 as compared to State 2 (Fig. 1A and (Chakraborty et al., 2020). Lineages were ranked (Figure 4A) and separated into 10 groups based on their **overall motility** (sum of motilities in all 4 transitions). The motility in each transition of each group is shown in Figures 4B, 4C, 4D and S4A. Percent Motility (used in Figure 4 and Figure S6) is calculated by:

**Percent Motility** = (Motility of that lineage / $\sqrt{2}$) * 100

**Calculating the correlation of motility among replicate experiments**

To determine whether the correlation in motility between two experimental replicates was significant (Figure S7C-D), we calculated a correlation of **randomized motility** using the following method: 1. Identify lineages that exist in both replicates; 2. For each replicate (WT1 and WT2), collect the experimental motility values of each transition for each lineage; 3. Randomly assigned motility values to each lineage from the distribution of motilities within that replicate using the data collected from step (2); 4. Calculate log_10_ of percent motilities from these randomly assigned WT1 and WT2 motility values; 5. For each lineage in WT1 and WT2 replicates, plot the data and calculate Pearson correlation coefficients (r-values) for each transition. The comparison between empirically observed motility correlation (Fig. S7C) and those randomly selected (Fig. S7D) is shown. Upon 100,000 trials of this randomization procedure, R^2^ between randomly assigned WT1 and WT2 motilities ranged from 0 to 0.01 for these trials.

**Fisher’s exact test of independence between different state histories**

To determine whether the difference in state distributions between two different state histories was significant (Supplemental Table 1, “Motif Analysis” sheet; Supplemental Table 2, “Motif Analysis” sheet), we compared the number of lineages in each state at Day 24 for lineages with two distinct histories for lineages that occupied the same state on Day 18 using Fisher’s Exact Test (fisher.test function in R). We restricted analysis to histories that contained lineages occupying all states in the last timepoint such that the Fisher statistic is defined. The contingency tables, p-values, and the Bonferroni corrected alpha value of *Nanog-Sox2* state transitions are found in Supplemental Table 1, “Fisher exact test” sheet. The contingency tables, p-values, and the Bonferroni corrected alpha value of *motility* state transitions are found in Supplemental Table 2, “Fisher exact test” sheet.
**Note**

**Estimating Transitional Probability Matrix**

We can model the system of three cell states where cells in each state can either stay in the same state or convert to one of the other two states after each transition. In this section, we describe how we estimate the transitional probabilities that explain how lineages change states between timepoints.

Each lineage contains information about the proportion of cells in different states, which is known for all timepoints (Day 0, Day 6, Day 12, Day 18 and Day 24). We will focus on two contiguous timepoints for clarity (note this will generalize to any transition): the data from Day 0 and Day 6.

Let $\mathbf{X}_{i}$ and $\mathbf{Y}_{i}$ be $1\times3$ row vectors that contain the proportion of 3 states of lineage $i$ on Day 0 and Day 6, respectively. In other words,

$$\begin{matrix} \mathbf{X}_{i} & =\left[ \begin{matrix} P_{s1,i} & P_{s2,i} & P_{s3,i} \end{matrix} \right] \\ \mathbf{Y}_{i} & =\left[ \begin{matrix} P'_{s1,i} & P'_{s2,i} & P'_{s3,i} \end{matrix} \right] \end{matrix}$$

Where for example $P_{s2,i}$ indicates the proportion of lineage i that is in State 2 at timepoint Day 0. Because these three states can interconvert between one another, there must be a matrix $\mathbf{M}$ of transitional probabilities can explain the conversion from $\mathbf{X}_{i}$ to $\mathbf{Y}_{i}$. Therefore, we can write,

$$\begin{matrix} \mathbf{X}_{i}\mathbf{M} & =\mathbf{Y}_{i} \end{matrix}$$

where

$$\begin{matrix} \mathbf{M} & =\left[ \begin{matrix} M_{11} & M_{12} & M_{13} \\ M_{21} & M_{22} & M_{23} \\ M_{31} & M_{32} & M_{33} \end{matrix} \right] \end{matrix}$$

Please note that the multiplication of $\mathbf{X}_{i}$ and $\mathbf{M}$ describes how state proportions $\left[ \begin{matrix} P_{s1,i} & P_{s2,i} & P_{s3,i} \end{matrix} \right]$ change to $\left[ \begin{matrix} P'_{s1,i} & P'_{s2,i} & P'_{s3,i} \end{matrix} \right]$.

Here, we want to solve a system of linear equations for matrix **M**. However, considering only one lineage leaves 3 constraints with 9 unknowns, making it difficult to explicitly consider $\mathbf{M}$ for any single lineage. Hence, we utilize the 2,560 lineages that exist in all timepoints (Fig. 1D). We can use this information to estimate the transitional probability matrix $\mathbf{M}$ that explains the transitions between states for **all** lineages on average between Day 0 to Day 6.

This now gives two $2560\times3$ matrices instead of two $1\times3$ row vectors. We note the matrix of state proportions of all lineages on Day 0 as $\mathbf{X}$ and its Day 6 counterpart as $\mathbf{Y}$. Note that row $j$ of $\mathbf{X}$ and $\mathbf{Y}$ represents the state proportions of lineage $j$ on Day 0 and Day 6, respectively. From this, we can write,

$$\begin{matrix} \mathbf{XM} & =\mathbf{Y} \end{matrix}$$

and can solve for the matrix $\mathbf{M}$ that minimizes the error of transforming $\mathbf{X}$ to $\mathbf{Y}$. Because this is a system of linear equations, we use ordinary least squares (OLS) estimation to calculate the matrix **M** by minimizing the sum of squared errors. The error of estimation is therefore given by $\mathbf{r}=\mathbf{Y-XT}$ where $\mathbf{r}$ is a 2560 × 3 matrix. We calculate the sum of squared errors (noted as $\mathbf{S}$) by multiplication of $\mathbf{r}$ with $\mathbf{r}^{T}$, where $\mathbf{r}^{T}$ denotes the transpose of $\mathbf{r}$.

$$\begin{matrix} \mathbf{S} & =<\mathbf{r},\mathbf{r}> \\ & =\mathbf{r}\mathbf{r}^{T} \\ & =(\mathbf{Y}-\mathbf{XM})(\mathbf{Y}-\mathbf{XM})^{T} \\ & =(\mathbf{Y}-\mathbf{XM})(\mathbf{Y}^{T}-\mathbf{M}^{T}\mathbf{X}^{T}) \\ & =\mathbf{Y}\mathbf{Y}^{T}-\mathbf{Y}\mathbf{M}^{T}\mathbf{X}^{T}-\mathbf{XM}\mathbf{Y}^{T}+\mathbf{XM}\mathbf{M}^{T}\mathbf{X}^{T} \end{matrix}$$

A matrix $\mathbf{M}$ that minimize the sum of squared errors $\mathbf{S}$ must occur at the critical point where the gradient of $\mathbf{S}$ with respect to $\mathbf{M}$ equals 0.

$$\begin{matrix} \frac{\partial\mathbf{S}}{\partial\mathbf{M}} & =\frac{\partial}{\partial\mathbf{M}}(\mathbf{Y}\mathbf{Y}^{T}-\mathbf{Y}\mathbf{M}^{T}\mathbf{X}^{T}-\mathbf{XM}\mathbf{Y}^{T}+\mathbf{XM}\mathbf{M}^{T}\mathbf{X}^{T}) \\ & =-2\mathbf{X}^{T}(\mathbf{Y}-\mathbf{XM}) \\ & =\mathbf{X}^{T}\mathbf{Y}-\mathbf{X}^{T}\mathbf{XM} \\ & =0 \end{matrix}$$

Therefore,

$$\begin{matrix} \mathbf{M} & =(\mathbf{X}^{T}\mathbf{X})^{-1}\mathbf{X}^{T}\mathbf{Y} \end{matrix}$$

Using the knowledge from this derivation, we can estimate the transitional probability matrix $\mathbf{M}$ that explains the transition from one timepoint to another subsequent timepoint by substituting $\mathbf{X}$ as a matrix of state proportions in an earlier timepoint and $\mathbf{Y}$ as a matrix of state proportions in a later timepoint.

In order to find an average transitional probability matrix (Fig. 3A) across all transitions, instead of using $2560\times3$ $\mathbf{X}$ and $\mathbf{Y}$ matrices, we combine the state proportions of all lineages on Day 0, Day 6, Day 12 and Day 18 into a $10240\times3$ matrix $\mathbf{X}'$ and combine the state proportions of all lineages on Day 6, Day 12, Day 18, and Day 24 into a $10240\times3$ matrix $\mathbf{Y'}$. Note that row $j$ of both $\mathbf{X'}$ and $\mathbf{Y'}$ represents the state proportions of the same lineage; $\mathbf{X'}_{j}$ has the information from the earlier timepoint while $\mathbf{Y'}_{j}$ has the information from the later timepoint. The results of this matrix are given (Fig. 3A).

**Estimating Growth-Birth-Death Rate**

We next sought to use our knowledge of transitional probabilities in the system of ESC lineages to estimate the net rate of growth, birth, and death events for mouse embryonic stem cells making all types of state transitions. Calculating the net growth-birth-death rate is not trivial because this rate is intertwined with state transition rates calculated in Section 1.1 and shown in Fig. 3A. In this section, we focus on extracting the net growth-birth-death rate once average rates of transition between states are taken into account.

We first find the distributions of cells in State 1, State 2 and State 3 across all lineages predicted by the transition matrix **M** for a given lineage making a transition between two timepoints. The number of cells of each lineage in each state at each timepoint is calculated from the empirical data as described (see Methods). We consider the difference between the number of cells at the second timepoint predicted by the transition matrix **M** and the number of cells empirically observed in each state at this timepoint for this lineage as a result of growth, birth, and death events.

To demonstrate, we will use the data from Day 0 and Day 6 as an example, noting that this procedure generalizes to analysis of data from all other contiguous timepoints. Let $\mathbf{U}_{i}$ and $\mathbf{V}_{i}$ be $1\times3$ row vectors that contain the normalized cell number in all three states for lineage $i$ on Day 0 and Day 6, respectively. In other words,

$$\begin{matrix} \mathbf{U}_{i} & =\left[ \begin{matrix} U_{s1,i} & U_{s2,i} & U_{s3,i} \end{matrix} \right] \\ \mathbf{V}_{i} & =\left[ \begin{matrix} V_{s1,i} & V_{s2,i} & V_{s3,i} \end{matrix} \right] \end{matrix}$$

The expected distribution of cells after Day 0 cells transition between states can be found by multiplying $\mathbf{U}_{i}$ with transitional probability matrix $\mathbf{M}$ we calculated in Section 1.1. The product is $\mathbf{U}'_{i}$.

$$\begin{matrix} \mathbf{U}'_{i} & =\mathbf{U}_{i} \mathbf{M} \end{matrix}$$

Here, vector $\mathbf{U'}_{i}$ describes the expected number of cells on Day 6 due to transition alone. Vector $\mathbf{V}_{i}$ contains the observed number of cells for lineage i on Day 6. We assume the difference between these two vectors stems from growth, birth, and death processes happening between Day 0 and Day 6 timepoints for this lineage. To model this process, we introduce a $3\times3$ matrix $\mathbf{G}$ which describes the difference between the number of cells in $\mathbf{U'}_{i}$ and $\mathbf{V}_{i}$. Mathematically,

$$\begin{matrix} \mathbf{U}'_{i} \mathbf{G} & =\mathbf{V}_{i} \end{matrix}$$

However, we cannot solve for matrix $\mathbf{G}$ for an individual lineage because the system of linear equations has more unknowns (9) than constraints (3). Hence, we again utilize the 2,560 lineages that are present at all timepoints (Fig. 1D), and use this information from all lineages to solve for $\mathbf{G}$ that minimizes the difference of state distributions using the ordinary least squares method.

Let $\mathbf{U}$ and $\mathbf{V}$ be $2560\times3$ matrices that represent the distributions of cells in three states of all 2,560 lineages on Day 0 and Day 6, respectively. We can write,

$$\begin{matrix} \mathbf{UG} & =\mathbf{V} \end{matrix}$$

From ordinary least square estimation,

$$\begin{matrix} \mathbf{G} & =(\mathbf{U}^{T}\mathbf{U})^{-1}(\mathbf{U}^{T}\mathbf{V}) \end{matrix}$$

Here $\mathbf{G}$ is a growth-birth-death transition matrix between Day 0 and Day 6. We can use this knowledge to find growth-birth-death transition matrices between two other contiguous timepoints.

We then use matrix $\mathbf{G}$ to find the expected number of cells in different states after the growth, birth and death processes and calculate the growth-birth-death rate for each cell state transition event.

Let a $1\times3$ row vector $\mathbf{N}$ describes the number of cells in different states. In other word, $\mathbf{N}=\left[ \begin{matrix} N_{s1} & N_{s2} & N_{s3} \end{matrix} \right]$. Assuming that there are 100 cells in State 1 at the first timepoint, we can find the expected number of cells $E(\mathbf{N})$ in different states after cell state transitions and growth-birth-death processes by calculating

$$\begin{matrix} \mathbf{N}_{s1} & =\left[ \begin{matrix} 100 & 0 & 0 \end{matrix} \right] \\ E(\mathbf{N}_{s1}) & =\mathbf{U}_{s1}\mathbf{MG} \end{matrix}$$

where $\mathbf{M}$ is the transitional probability matrix we derived from the least square estimation and $\mathbf{G}$ is the growth-birth-death transitions derived above.

The rate of change in State 1 number due to the growth, birth, death processes is simply the entrywise division (Hadamard division) between $E(\mathbf{N}_{s1})$ and $\mathbf{N}_{s1}\mathbf{M}$. In other words, Growth-birth-death rate${}_{s1}=\left[ \begin{matrix} E(\mathbf{N}_{s1})_{11}/(\mathbf{N}_{s1}\mathbf{M})_{11} & E(\mathbf{N}_{s1})_{12}/(\mathbf{N}_{s1}\mathbf{M})_{12} & E(\mathbf{N}_{s1})_{13}/(\mathbf{N}_{s1}\mathbf{M})_{13} \end{matrix} \right]$ where $E(\mathbf{N}_{s1})_{ij}$ is the element $i,j$ in $E(\mathbf{N}_{s1})$ and ($\mathbf{N}_{s1}\mathbf{M})_{ij}$ is the element $i,j$ in $\mathbf{N}_{s1}\mathbf{M}$.

We can calculate the rate of change in the number of cells in State 2 and 3 using the method above.

To calculate an overall growth-birth-death rate across all 5 timepoints, we form a matrix $\mathbf{U'}$ that contains the number of cells in each state from all lineages from 4 timepoints (Day 0, Day 6, Day 12 and Day 18) and form a matrix $\mathbf{V'}$ that contains the number of that contains the number of cells in each state from all lineages from 4 subsequent timepoints (Day 6, Day 12, Day 18 and Day 24). Please note that entries in each row of $\mathbf{U'}$ and $\mathbf{V'}$ are from the same lineages; entries in $\mathbf{U'}$ are from an earlier timepoint while their counterparts in $\mathbf{V'}$ are from the subsequent timepoint. From this we can calculate the growth-birth-death transition matrix $\mathbf{G'}$ and the growth-birth-death rate for each state the way we have mentioned above. Rates inferred from G’ are shown in Fig. S3A.

**Calculating Lineage Entropy**

We start here with definitions of concepts as they are considered in the present study; these may be familiar to many readers. In information theory, entropy of a variable reveals the average amount of information or uncertainty in its outcomes. Given a random variable X with n possible outcomes $x_{1},x_{2},...,x_{n}$ that occurs with probability $P(x_{1}),P(x_{2}),...,P(x_{n})$, the informational entropy of $X$ can be mathematically defined as

$$\begin{matrix} H(X) & =-\sum_{i=1}^{n} P(x_{i})\text{log}P(x_{i}) \end{matrix}$$

where entropy is always between 0 and 1. This quantity may be familiar to readers as the Shannon Entropy.

A further intuition can be found by considering a coin-flip thought experiment. If the coin is fair, there are two possible outcomes (heads and tails) both occuring at equal probability 1/2. Therefore, if flipping this coin, we do **not** know for sure which result will be obtained, meaning this system has maximal uncertainty and maximal information to be gained once we know the result of the coin flip, high information entropy). The entropy in this system in this case is

$$\begin{matrix} H(X) & =-\sum_{i=1}^{n} P(x_{i})\text{log}P(x_{i}) \\ & =-(\frac{1}{2}\text{log}\frac{1}{2}+\frac{1}{2}\text{log}\frac{1}{2}) \\ & =1 \end{matrix}$$

On the other hand, if we are flipping a coin that has heads on both sides, we do know for sure that after the coin flip the result will be heads and this does not yield us any new information (minimal uncertainty, minimal informational entropy, minimal information to be gained once the result of the coin flip is known). Mathematically,

$$\begin{matrix} H(X) & =-\sum_{i=1}^{n} P(x_{i})\text{log}P(x_{i}) \\ & =-(1\text{log}1) \\ & =0 \end{matrix}$$

In general, the system with more uncertainty is considered to have more information content to be gained and higher informational entropy value.

We can use this concept to describe the lineage entropy (or informational entropy) in each lineage, consistent with the idea that lineages with more heterogeneous proportion amongst states should have higher entropy than lineages occupying only one cell state, in analogy to our coin flip.

According to the empirical data, the steady-state distribution of State 1, State 2 and State 3 given by averaging all values is at $\left[ \begin{matrix} P(s1) & P(s2) & P(s3) \end{matrix} \right]=\left[ \begin{matrix} 0.756 & 0.126 & 0.118 \end{matrix} \right]$. We define the lineage with these specific proportions to have maximum lineage entropy equal to 1. However, this means we must scale state distributions when using a function of the form given by the Shannon entropy equation above (otherwise the maximum would occur at $\left[ \begin{matrix} \frac{1}{3} & \frac{1}{3} & \frac{1}{3} \end{matrix} \right]$, which is not the average steady state of the ESC system). We calculate the lineage entropy H’(L) of a lineage L by:

$$\begin{matrix} H'(L) & =-(P'(s1)\text{log}P'(s1)+P'(s2)\text{log}P'(s2)+P'(s3)\text{log}P'(s3)) \end{matrix}$$

where $P'(s1),P'(s2)$ and $P'(s3)$ are the scaled proportion of State 1, State 2 and State 3 in lineage $L$ respectively. In other words,

$$\begin{matrix} P'(s1) & =\left\{ \begin{matrix} \frac{P(s1)}{0.756}\times\frac{1}{3} & 0\leq P(s1)\leq0.756 \\ 1+\frac{P(s1)-1}{1-0.756}\times\frac{2}{3} & 0.756<P(s1)\leq1 \end{matrix} \right. \\ P'(s2) & =\left\{ \begin{matrix} \frac{P(s2)}{0.126}\times\frac{1}{3} & 0\leq P(s2)\leq0.126 \\ 1+\frac{P(s2)-1}{1-0.126}\times\frac{2}{3} & 0.126<P(s2)\leq1 \end{matrix} \right. \\ P'(s3) & =\left\{ \begin{matrix} \frac{P(s3)}{0.118}\times\frac{1}{3} & 0\leq P(s3)\leq0.118 \\ 1+\frac{P(s3)-1}{1-0.118}\times\frac{2}{3} & 0.118<P(s3)\leq1 \end{matrix} \right. \end{matrix}$$

This lineage entropy is calculated and shown in Figure 3D-F.

Bhang, H.E., Ruddy, D.A., Krishnamurthy Radhakrishna, V., Caushi, J.X., Zhao, R., Hims, M.M., Singh, A.P., Kao, I., Rakiec, D., Shaw, P.*, et al.* (2015). Studying clonal dynamics in response to cancer therapy using high-complexity barcoding. Nat Med *21*, 440-448.

Chakraborty, M., Hu, S., Visness, E., Del Giudice, M., De Martino, A., Bosia, C., Sharp, P.A., and Garg, S. (2020). MicroRNAs organize intrinsic variation into stem cell states. Proc Natl Acad Sci U S A *117*, 6942-6950.
